# Genome-wide screening reveals metabolic regulation of translational fidelity

**DOI:** 10.1101/2022.10.26.513850

**Authors:** Zhihui Lyu, Patricia Villanueva, Liam O’Malley, Parker Murphy, Jiqiang Ling

## Abstract

Translational quality control is critical for maintaining the accuracy of protein synthesis in all domains of life. Mutations in aminoacyl-tRNA synthetases and the ribosome are known to affect translational fidelity and alter fitness, viability, stress responses, neuron function, and life span. In this study, we used a high-throughput fluorescence-based assay to screen a knock-out library of *Escherichia coli* and identified 30 nonessential genes that are critical for maintaining the fidelity of stop-codon readthrough. Most of these identified genes have not been shown to affect translational fidelity previously. Intriguingly, we show that several genes controlling metabolism, including *cyaA* and *guaA*, unexpectedly enhance stop-codon readthrough. CyaA and GuaA catalyze the synthesis of cyclic adenosine monophosphate (cAMP) and guanosine monophosphate (GMP), respectively. Both CyaA and GuaA increase the expression of ribosomes and tRNAs, allowing aminoacyl-tRNAs to compete with release factors and suppress stop codons. In addition, the effect of *guaA* deletion on stop-codon readthrough is abolished by deleting *prfC*, which encodes release factor 3 (RF3). Our results suggest that nucleotide and carbon metabolism is tightly coupled with translational fidelity.

## INTRODUCTION

Accurate translation of message RNAs (mRNA) into proteins is a central and delicately controlled cellular process (1–3). First, aminoacyl-tRNA synthetases (aaRSs) are challenged to ligate the correct amino acids to the tRNAs (4). Given the similarity between amino acids, an editing step is often required to remove misactivated amino acids or mismatched aminoacyl-tRNAs (aa-tRNAs) (2,5,6). Second, the accurate pairing of the resulting aa-tRNAs to the mRNA codons on the ribosome also necessitates kinetic proofreading (7). Despite such conserved quality control mechanisms, base levels of translational errors still remain. The amino acid misincorporation rate at sense codons is approximately 10^−4^ to 10^−3^ and the frequency of stop-codon readthrough is even higher at 10^−3^ to 10^−2^ (8–12). It is important to note that such error rates are not fixed and can fluctuate due to mutations, mRNA context, and environmental changes (2,13–17). Mutations in aaRSs and tRNAs may lead to the production of misacylated tRNAs, which are used by the ribosome to make erroneous proteins (6,18–22). Recent studies suggest that some tRNA variants in human populations lead to increased translational errors and could function as disease modifiers (17,23). Similarly, mutations in ribosomal proteins and RNAs can also affect the decoding accuracy and cause amino acid misincorporation, stop-codon readthrough, and frameshift (11,24,25). In addition, programmed stop-codon readthrough and frameshift are frequently used by viruses and host cells to regulate the expression of alternative protein variants (26,27). In addition to genetic changes, it has also been shown that environmental cues can affect various types of translational errors. For example, oxidative stress impairs the editing function of threonyl-tRNA synthetase and causes serine misincorporation at threonine codons (28). Conversely, oxidation of phenylalanyl-tRNA synthetase enhances editing and confers hyper accuracy (16). Amino acid starvation and acid stress have also been shown to increase stop-codon readthrough on the ribosome (14,29).

Reduced translational fidelity often leads to detrimental effects such as slow growth, cell death, and neurological disorders (30). Recent studies in yeasts, flies, worms, and mice also demonstrate that increasing translational fidelity by a ribosomal mutation improves the life span, whereas decreasing translational fidelity leads to accelerated aging (31,32). However, maintaining high fidelity also comes with costs, and increasing translational errors have been shown to be beneficial under certain conditions. For example, increased ribosomal decoding fidelity decreases resistance to oxidative stress in *Escherichia coli* and the expression of virulence genes in *Salmonella enterica*, suggesting that the base level of translational errors improves bacterial fitness (33,34). Elevating specific types of translational errors also increase resistance against antibiotics in *Mycobacteria* and against oxidative stress in mammalian cells (35–37).

To understand what genetic factors and pathways contribute to the regulation of translational fidelity, we used a recently developed dual-fluorescence reporter to perform high-throughput screening of the *E. coli* knockout library. In addition to a few genes that were previously shown to affect translational fidelity (e.g., *miaA* and *prfC*), we identified over 20 novel genes that either increase or decrease the readthrough of UGA stop codons. We performed validation by deleting these genes in an independent *E. coli* strain MG1655. Our results further revealed that several metabolic genes affected stop-codon readthrough by regulating the activities of translational factors and the biogenesis of tRNAs and ribosomes.

## RESULTS

### High-throughput screening for *E. coli* knockout mutants with altered UGA readthrough

Increasing evidence shows that translational fidelity is regulated by genetic and environmental cues, and plays crucial roles in fitness and adaptation. We have previously constructed dual-fluorescence reporters to detect stop-codon readthrough in populations and single cells (12). These reporters allow convenient and high-throughput quantitation of translational errors. To search for novel genes that are important for translational fidelity, we introduced the UGA readthrough reporter (pZS-Ptet-m-TGA-y) and the control (pZS-Ptet-m-y) plasmids into the Keio knockout collection (38) in 96-well plates (Figure 1). The transformants were then cultured in fresh Lennox broth (LB) over 16 hours at 25 and 37 °C, respectively. The mCherry and YFP signals were quantitated with a platereader, and the UGA readthrough levels were determined using the ratio of YFP over mCherry as described (12).

**Figure 1.**
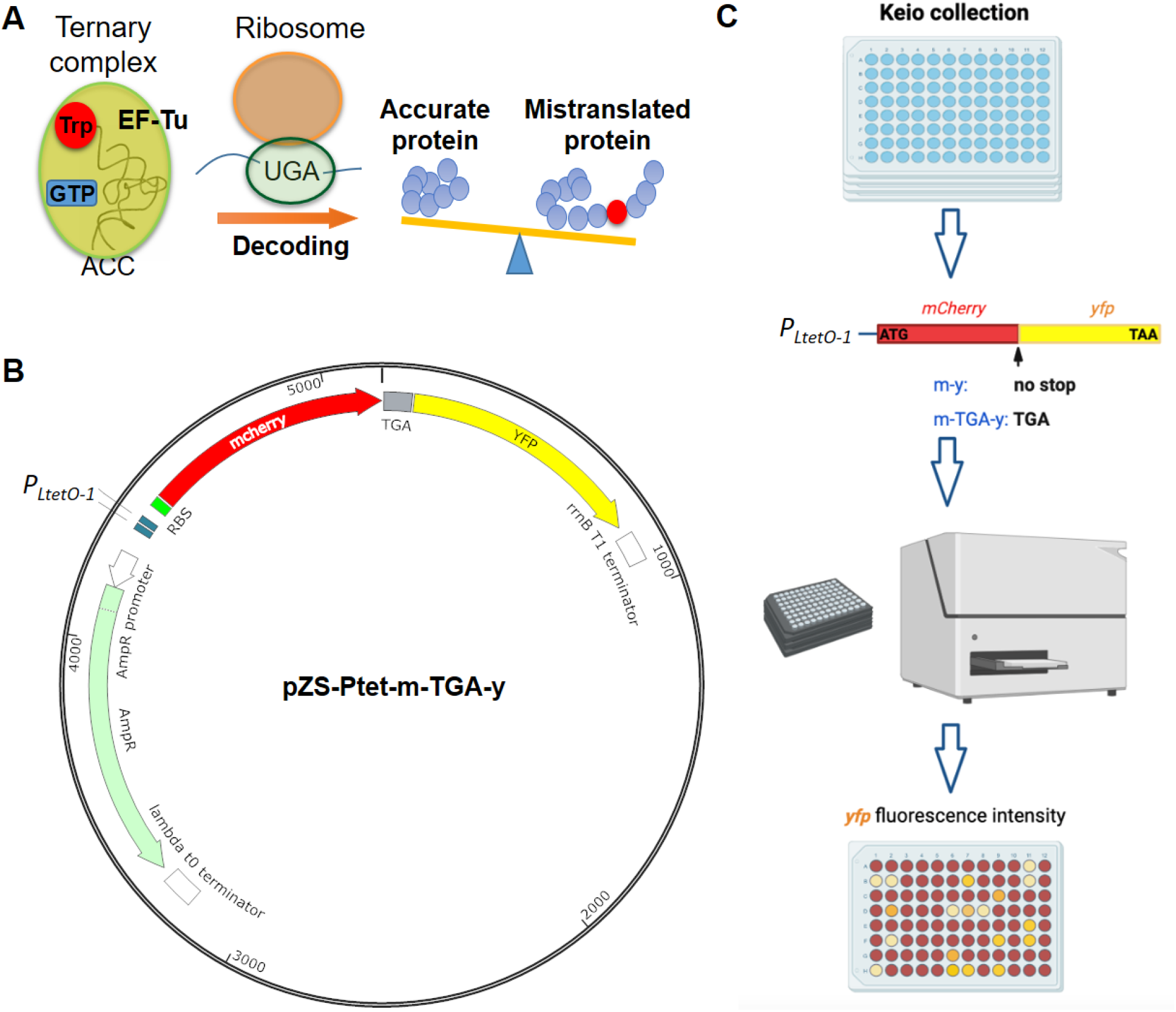
Genome-wide screening of *E. coli* knockout library for altered UGA readthrough levels. **(A)** The EF-Tu:Trp-tRNA^Trp^:GTP ternary complex competes with release factors to suppress the UGA stop codon and results in extended proteins. **(B)** Map of the UGA readthrough reporter plasmid. YFP, yellow fluorescence protein. **(C)** Workflow of high-throughput screening for *E. coli* knockout mutants with altered UGA readthrough. The plasmids were transformed into the Keio knockout collection using 96-well plates. The transformants were then cultured and measured for fluorescence in a platereader.

A total of ~ 4000 strains covering the entire Keio collection were screened. Growing the wild-type strain BW25113 at 25 and 37 °C yielded ~ 4% and 2% of UGA readthrough, respectively (Figure 2). Using the cutoff of either ≥ 50% increase or ≥ 33% decrease in the UGA readthrough level at 25 °C, we identified a total of 30 knockout strains. Based on the functions, we grouped the 30 genes into metabolism, translation, transcription, redox, and other (Figure 2 and Table S1). Among the identified genes from our screening, *miaA, prfC, rplI*, and *rsmG* are translational factors that have been previously associated with translational fidelity. MiaA catalyzes the isopentenyladenosine modification at position 37 (i^6^A37) of tRNAs and its deletion decreases UGA readthrough (39); it also plays an important role in the general stress response (40). The *prfC* gene encodes RF3, which facilitates RF1 and RF2 to terminate translation at stop codons (41). *rplI* encodes the L9 protein of the ribosome and deleting *rplI* increases translational readthrough, frameshift, and recoding of stalled ribosomes (42,43). RsmG (GidB) is a ribosomal RNA (rRNA) methyltransferase and has been shown to increase some types of amino acid misincorporation in *Mycobacteria tuberculosis* (44). Among the newly identified genetic factors that affect translational fidelity, *tufA* is one of the two genes that encode elongation factor Tu (EF-Tu). During translation termination, EF-Tu forms a ternary complex (TC) with aa-tRNA and GTP to compete with release factors and suppress stop codons (Figure 1A). A reduction in TC in the Δ*tufA* strain would explain a decrease in UGA readthrough (Figure 2). A large fraction of the remaining genes are involved in cellular metabolism. For instance, CyaA catalyzes the synthesis of cAMP, which regulates the expression of many metabolic pathways when bound to Crp (45); GuaA is one of the two enzymes in *E. coli* that synthesizes GMP (46); SpeA contributes to the biosynthesis of polyamines (47).

**Figure 2.**
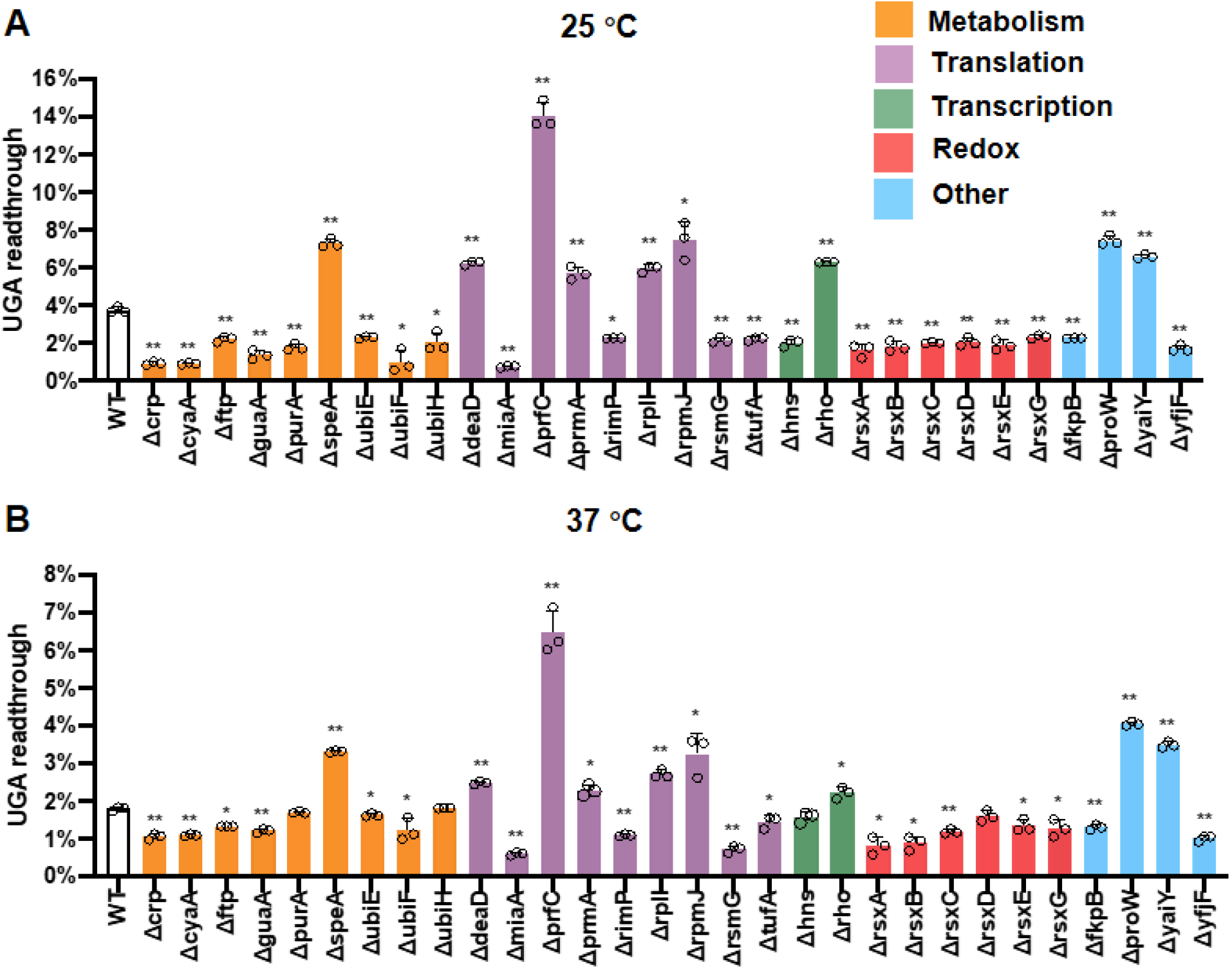
Genes affecting UGA readthrough revealed by screening. Overnight cultures of *E. coli* with the UGA readthrough reporter (pZS-Ptet-m-TGA-y) and the control (pZS-Ptet-m-y) were diluted 1:50 in fresh LB Amp over 16 hours at 25 **(A)** and 37 °C **(B)**, respectively. The fluorescence intensity of mCherry and YFP was determined using a platereader. UGA readthrough rates were calculated using the normalized ratio of YFP over mCherry. Error bars indicate standard deviations. P values were calculated using the unpaired t-test comparing the mutants with the WT. * P ≤ 0.05, ** P ≤ 0.0001.

To validate the roles of the identified genes in UGA readthrough, we performed complementation experiments by introducing ASKA overexpression plasmids into the corresponding knockout strains (48). UGA readthrough of most deletion strains was largely restored to the wild-type (WT) level, except for Δ*guaA*, Δ*prmA*, Δ*rimP*, Δ*rpmJ*, Δ*rho*, Δ*fkpB*, Δ*proW*, Δ*yaiY*, and Δ*yfjF* (Figure S1). We reasoned that the lack of complementation could result from secondary mutations in the Keio knockout strains, dosage effects of gene expression, polar effects on the expression of neighboring genes, or nonfunctional proteins due to the N-terminal histidine tag on the ASKA plasmids.

### Independent validation of translational errors using knockout mutants derived from MG1655

To rule out the effects of nonspecific mutations in the Keio strains on UGA readthrough, we constructed 18 deletion mutants from *E. coli* MG1655. The 12 other genes were left out due to the redundant pathways (*crp* and *rsxBCDEF*), small effect on UGA readthrough at 37 °C (*purA* and *hns*), difficulty in deletion (*ubiEFH*), or the lack of complementation (*rho*). The UGA readthrough levels of MG1655-derived mutants all confirmed the changes seen in the Keio strains except for Δ*rimP*, Δ*proW*, and Δ*yaiY*, which showed no significant change in UGA readthrough at 25 °C (Figure 3A).

**Figure 3.**
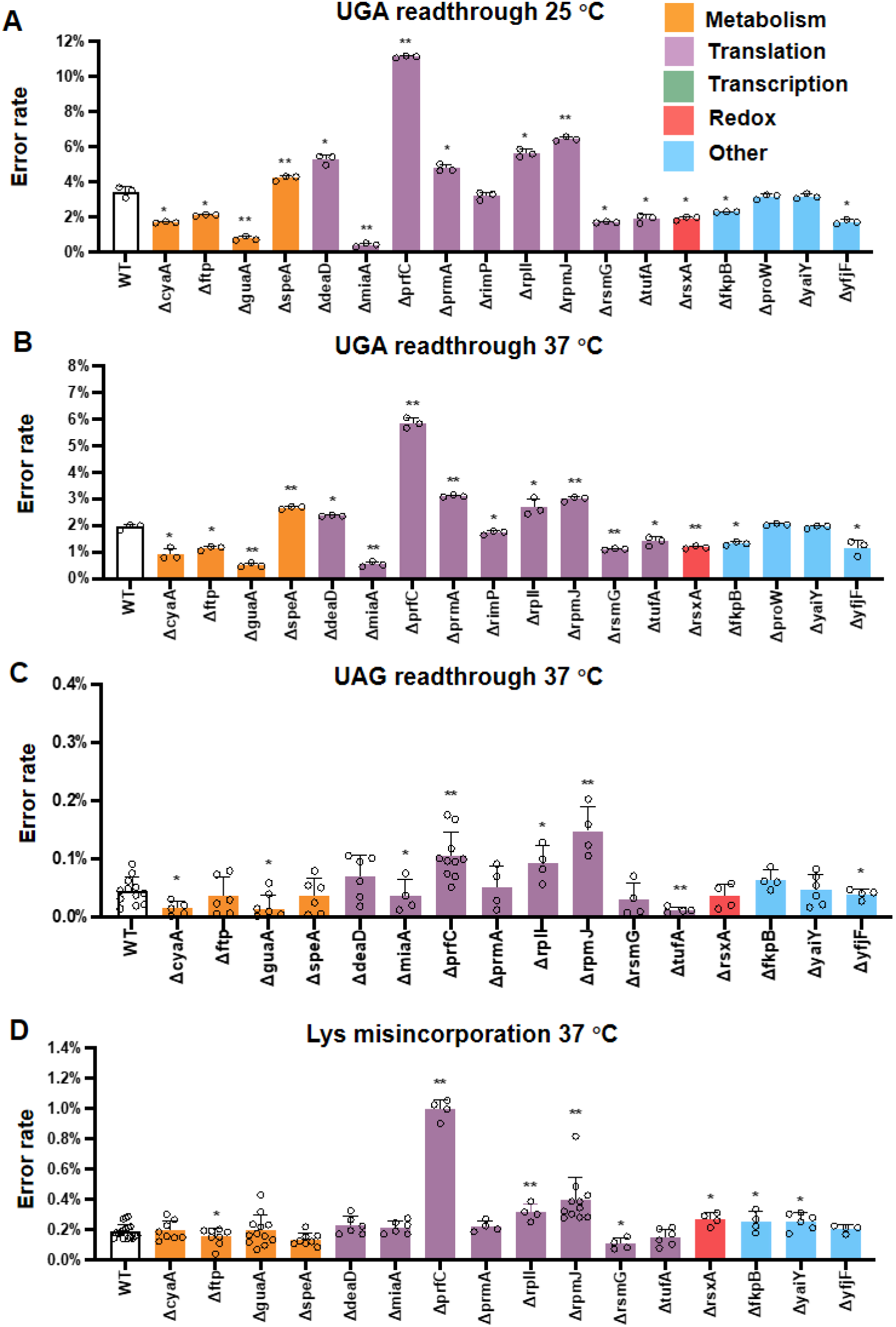
Error rates of mutants derived from MG1655. UGA readthrough reporter (pZS-Ptet-m-TGA-y) and the control (pZS-Ptet-m-y) plasmid were transformed into *E. coli* MG1655 mutant strains. Overnight cultures of *E. coli* were diluted 1:50 in LB Amp and incubated at 25 °C **(A)** or 37 °C **(B)** for 16 h. UAG readthrough (**C)** and lysine misincorporation **(D)** rates were determined using dual-luciferase reporters. Error bars indicate standard deviations. P values were calculated using the unpaired t-test comparing the mutants with the WT. * P ≤ 0.05, ** P ≤ 0.0001.

We next tested how the deletion of these genes affects other types of translational errors. Due to the low frequencies of UAG readthrough and misincorporation, we used dual-luciferase reporters as previously described (8,11). Deleting *cyaA, guaA, miaA, tufA*, and *yfjF* decreased both UAG and UGA readthrough but had no effect on lysine misincorporation at AAU asparagine codons (Figure 3). This suggests that these genes specifically influence stop-codon readthrough. On the other hand, deleting *prfC, rplI, rpmJ*, or *rsmG* affected both stop-codon readthrough and amino acid misincorporation, indicating their broader roles in the regulation of translational fidelity.

The growth rate of WT MG1655 was twice at 37 °C as compared to 25 °C (Figure S2), yet the UGA readthrough rate was about 50% lower at 37 °C than that at 25 °C (Figure 3). To test if growth rates are broadly associated with the UGA readthrough levels, we performed growth assays of MG1655-derived mutants and did not observe an apparent correlation between the two (Figure S2). In addition, UGA readthrough did not appear to correlate with the cellular ATP levels, either (Figure S3).

### Single-cell analyses of UGA readthrough in *E. coli* variants

Our dual-fluorescence reporter (m-TGA-y) not only empowers us to perform high-throughput screening but also allows us to observe UGA readthrough in single cells (12,14). We detected decent YFP signals resulting from readthrough in WT cells using both fluorescence microscopy (Figures 4 and S4) and flow cytometry (Figure S5). Overall, UGA readthrough was heterogeneous among single cells in the same population, and the single-cell results agreed nicely with the population-based platereader results (Figures 2–4, S4, and S5). Both Δ*cyaA* and Δ*guaA* cells showed brighter mCherry signals compared with the WT (Figure 4), presumably due to slow growth (Figure S2) that results in the accumulation of stable fluorescent proteins. Despite the high mCherry levels, the YFP signals in the Δ*cyaA* and Δ*guaA* cells were weaker than in the WT (Figure 4), suggesting lower UGA readthrough levels in these mutant cells. The Δ*guaA* cells also appeared to be longer than the WT, implying a possible defect in cell division (Figure 4).

**Figure 4.**
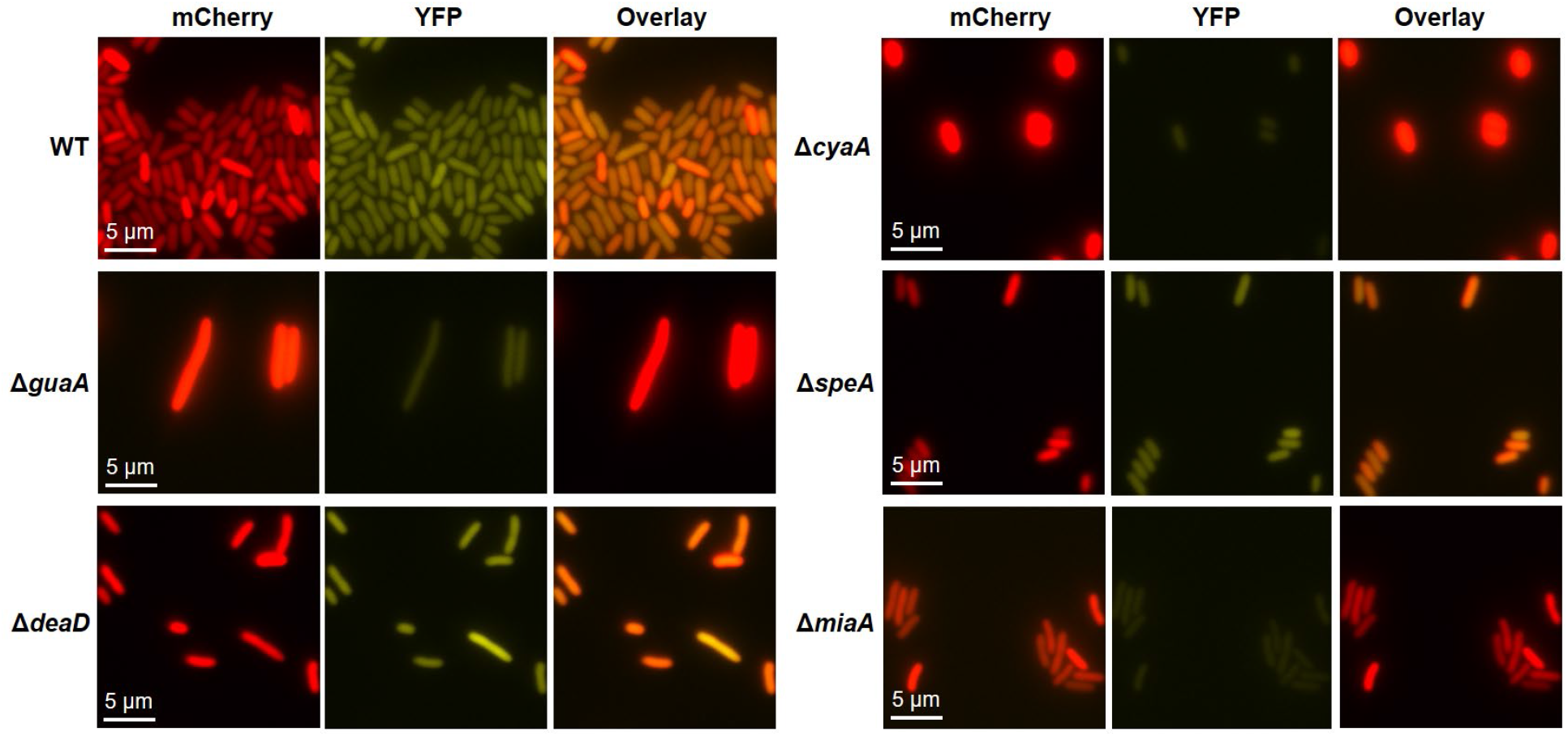
UGA readthrough in single cells. Overnight cultures of *E. coli* variants (derived from MG1655) carrying pZS-Ptet-mCherry-TGA-y were diluted 1:50 in LB Amp and grown aerobically at 25 °C for 16 h with shaking. Fluorescence images were obtained using BZ-X800 fluorescence microscope with a 100x oil lens.

### Impact of release factors on UGA readthrough

The UGA readthrough level is primarily determined by the competition between the EF-Tu:Trp-tRNA^Trp^:GTP ternary complex and RF2, which is facilitated by GTP-bound RF3 (12,41). Overexpressing RF2 or RF3 did not substantially decrease UGA readthrough in most of the tested variants (Figure 5A), suggesting that the release factor activity was largely saturated. As expected, overexpressing RF3 substantially decreased UGA readthrough in the Δ*prfC* mutant.

**Figure 5.**
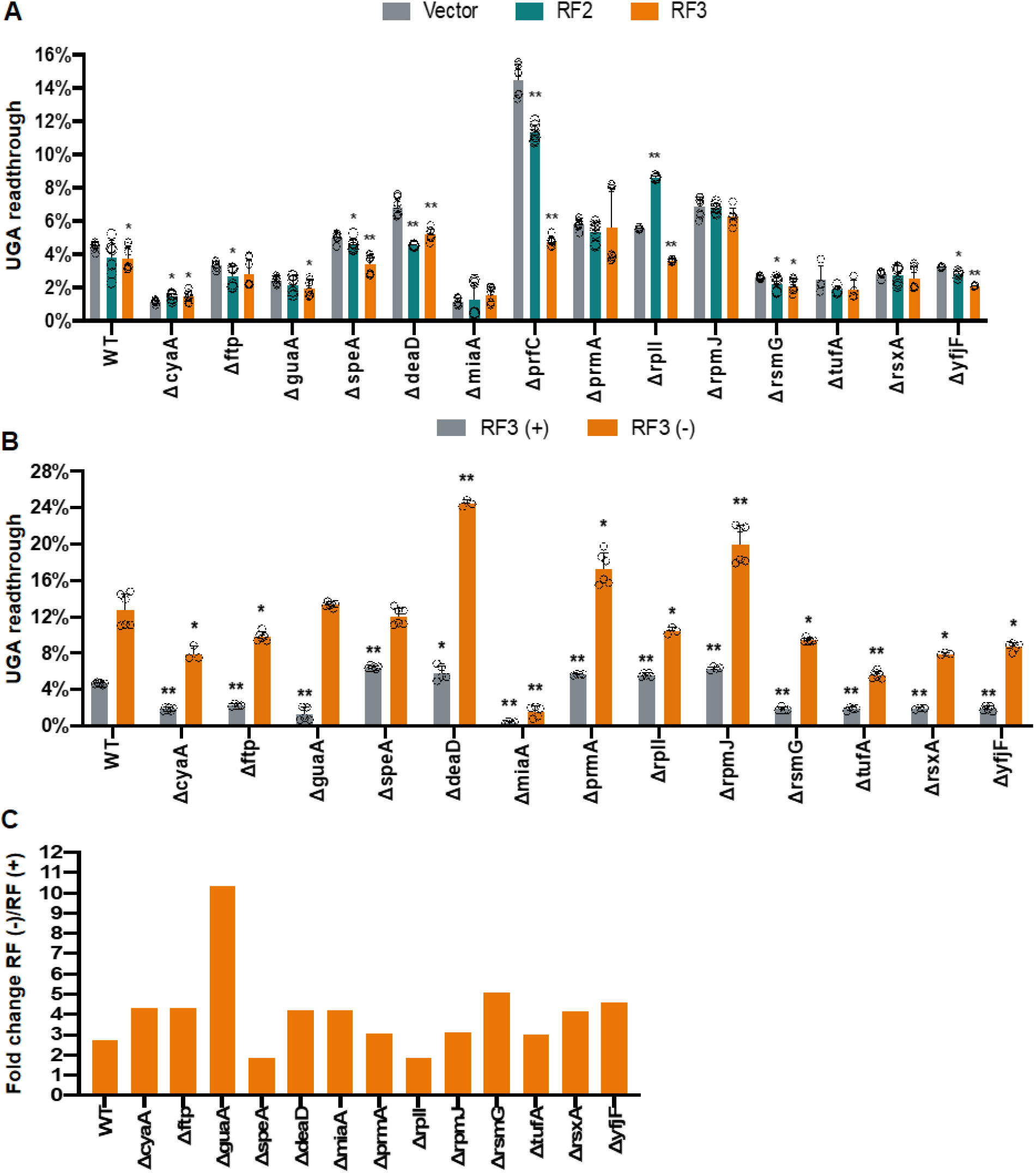
Effects of release factors on UGA readthrough in *E. coli* mutants. **(A)** ASKA plasmids expressing RF2 or RF3 were transformed into *E. coli* variants with pZS-Ptet-m-TGA-y and pZS-Ptet-m-y. The UGA readthrough levels were determined as in Figure 2. **(B**) and **(C)** UGA readthrough in the presence and absence of RF3. Error bars indicate standard deviations. P values were calculated using the unpaired t-test comparing the mutants with the WT carrying the same plasmids. * P ≤ 0.05, ** P ≤ 0.0001.

We have recently shown that the addition of glucose in rich media enhances stop-codon readthrough by reducing media pH, which impairs the activity of RF2 (14). Except for Δ*miaA*, all tested strains displayed an increase in UGA readthrough upon the addition of 1% glucose in LB (Figure S6). This suggests that tRNA^Trp^ lacking the i^6^A37 modification does not efficiently suppress the UGA codon even if the RF2 activity is impaired.

We next deleted RF3 in the WT and single knockout mutants. Further deletion of RF3 increased UGA readthrough in all the tested variants (Figures 5B and 5C). interestingly, deleting RF3 in Δ*guaA* and WT abolished the difference in UGA readthrough between the two strains, implying that GuaA primarily affects UGA readthrough via RF3. As GuaA facilitates biosynthesis of GMP and GTP, it likely enhances stop-codon readthrough by increasing the level of GTP-bound RF3.

### Increasing tRNA^Trp^ and tryptophan levels enhances UGA readthrough in *E. coli* variants

The TC/RF competition model for UGA readthrough indicates that fluctuation in tRNA^Trp^ and Trp levels may affect UGA readthrough (12). To test this, we overexpressed tRNA^Trp^ from a plasmid in the WT and mutant strains and observed a substantial increase in UGA readthrough in all strains (Figure S7). The level of tRNA^Trp^ was found to increase in the presence of chloramphenicol (Chl) (12). Indeed, the addition of Chl increased UGA readthrough in all the tested strains (Figure S6). Further, the addition of 40 mM Trp in LB also increased UGA readthrough (Figure S8). These data support that stop-codon readthrough is flexible in cells and can be affected by fluctuation in the levels of amino acids and tRNAs.

### Nucleotide metabolic genes *cyaA* and *guaA* enhance the biogenesis of ribosomes and tRNA^Trp^

Our screening revealed several metabolic genes to critically affect stop-codon readthrough. Our limited knowledge of how metabolism regulates translational fidelity prompted us to further test the roles of *cyaA, guaA*, and *speA* in regulating stop-codon readthrough. SpeA contributes to the biosynthesis of polyamines. Previous studies have shown that polyamines decrease the misincorporation of certain amino acids *in vitro* and *in vivo* (49,50). Here we found that the addition of 5 mM spermidine decreased UGA readthrough in both the WT and Δ*speA* mutant to the same level (Figure 6), suggesting that polyamines also contribute to the fidelity of translational termination.

**Figure 6.**
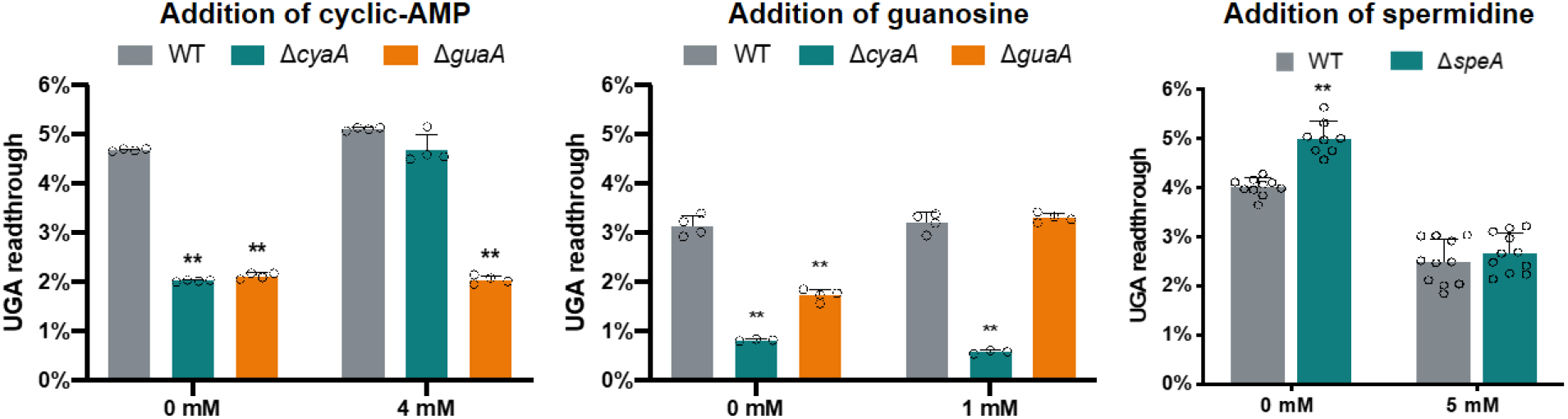
Metabolites restore UGA readthrough in Δ*speA*, Δ*cyaA*, and Δ*guaA* mutants. Overnight cultures of *E. coli* with pZS-Ptet-m-TGA-y and (pZS-Ptet-m-y were diluted 1:50 in LB Amp supplemented with mM cAMP, 1 mM guanosine, or 5 mM spermidine. The UGA readthrough levels were determined as in Figure 2. Error bars indicate standard deviations. P values were calculated using the unpaired t-test comparing the mutants with the WT. * P ≤ 0.05, ** P ≤ 0.0001.

CyaA and GuaA are both involved in nucleotide metabolism. CyaA is the only adenylate cyclase in *E. coli* to produce cAMP. Using an eCFP reporter under the control of the *tnaC* promoter, we confirmed that the CRP-cAMP activity was abolished in the absence of *cyaA* (Figure S9). The addition of 4 mM cAMP in the media fully restored *tnaC* transcription and UGA readthrough of the Δ*cyaA* mutant to the WT levels (Figures 6 and S9). In contrast, the addition of cAMP was not able to increase UGA readthrough in the Δ*guaA* strain. GuaA converts xanthosine monophosphate to GMP, which can be further used to synthesize GTP (46). In *E. coli*, a second pathway utilizes guanine or guanosine to synthesize GMP (46). We found that the addition of guanosine fully restored UGA readthrough to the WT level (Figure 6), suggesting that the decreased UGA readthrough in the Δ*guaA* strain is due to a GMP (and GTP) biosynthesis defect.

Many tRNAs are transcribed in the same operon as rRNAs. For instance, the gene encoding tRNA^Trp^ is part of the *rrsC* rRNA operon in *E. coli*. We speculated that dysregulation of nucleotides in the Δ*cyaA* and Δ*guaA* mutants may affect transcription of rRNAs and tRNAs. We thus determined the promoter activities of *rrsC* and the levels of tRNA^Trp^ in the WT, Δ*cyaA*, and Δ*guaA* variants (Figure 7). Both deletion strains exhibited a decrease in the tRNA^Trp^ level (Figures 7C and D). In line with this, Δ*cyaA* and Δ*guaA* also displayed a decreased RNA/protein ratio (Figure 7B), which was used to estimate the ribosome content in cells (51). The polysome profiles of the Δ*cyaA* and Δ*guaA* strains were similar to that of the WT except for a lower 70S peak (Figure S10), consistent with decreased ribosome biogenesis. Collectively, these data suggest that CyaA and GuaA are important in the biogenesis of rRNAs and tRNAs, and a decrease in the tRNA^Trp^ level contributes to the lower UGA readthrough activities observed in Δ*cyaA* and Δ*guaA* mutants.

CRP-cAMP is a central regulator of carbon metabolism (45). We reasoned that growth conditions with different carbon sources could affect UGA readthrough in the Δ*cyaA* strain. Indeed, when growing *E. coli* in minimal media (M9) with glucose as the carbon source, deleting *cyaA* caused higher UGA readthrough than the WT (Figure 8), the opposite of growth in LB with peptides as the carbon source (Figure 3). In contrast, all other deletions showed similar effects on UGA readthrough in M9 and LB. These results support that the regulation of stop-codon readthrough by CRP-cAMP depends on carbon metabolism.

**Figure 7.**
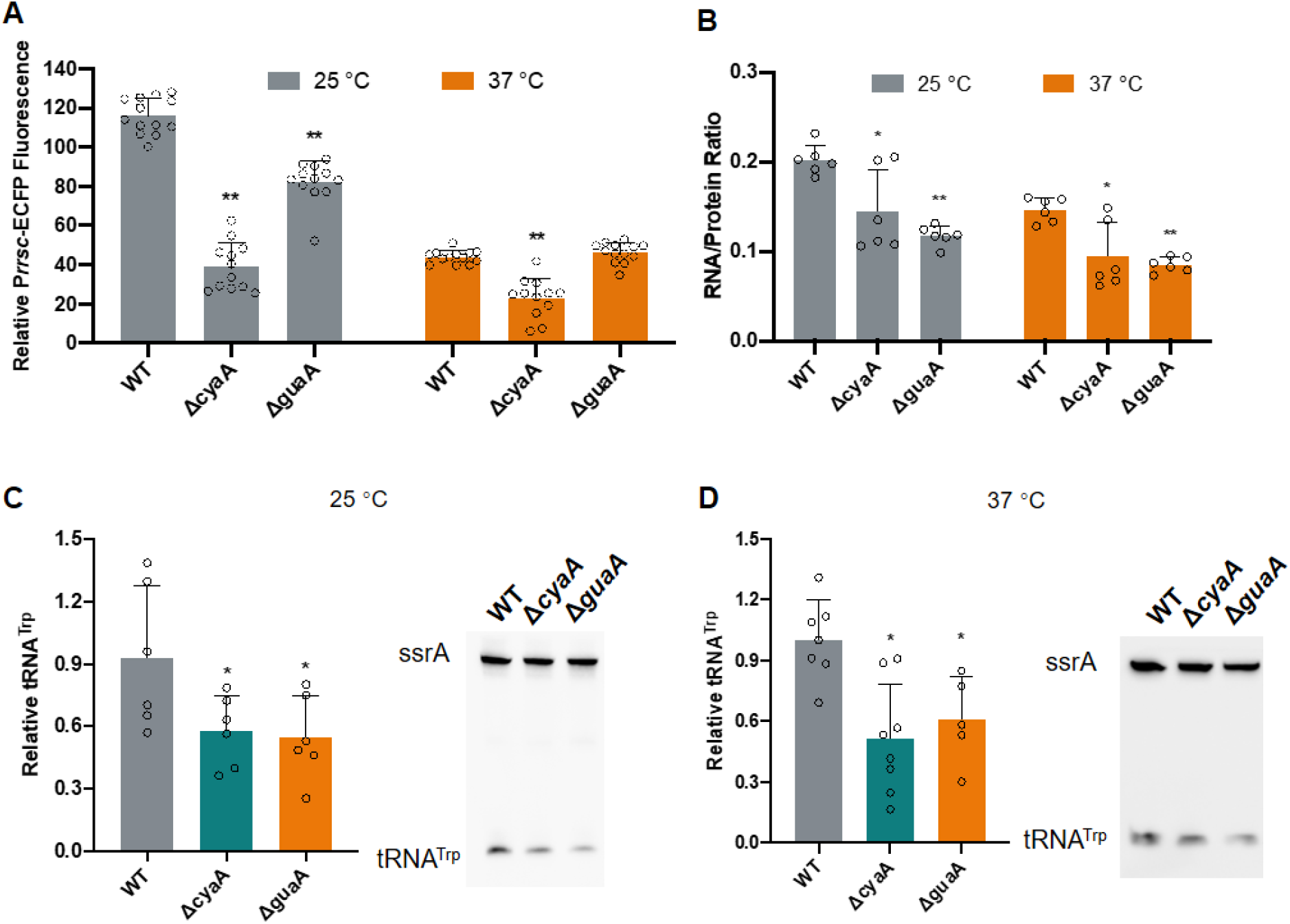
CyaA and GuaA regulate biogenesis of rRNA and tRNATrp. **(A)** Promoter activities of *rrsC*. Overnight cultures harboring pZS-m-TGA-y-P*rrsC*-eCFP were diluted 1:50 in fresh LB Amp and incubated at 25 °C or 37 °C for 16 h. The promoter activity was calculated as the ratio of eCFP over mCherry. **(B)** RNA/protein ratio. Cells grown at 25 °C or 37 °C to the mid-log phase in LB. **(C)** and **(D)** The levels of tRNA^Trp^ shown by northern blotting. Error bars indicate standard deviations. P values were calculated using the unpaired t-test comparing the mutants with the WT. * P ≤ 0.05, ** P ≤ 0.0001.

**Figure 8.**
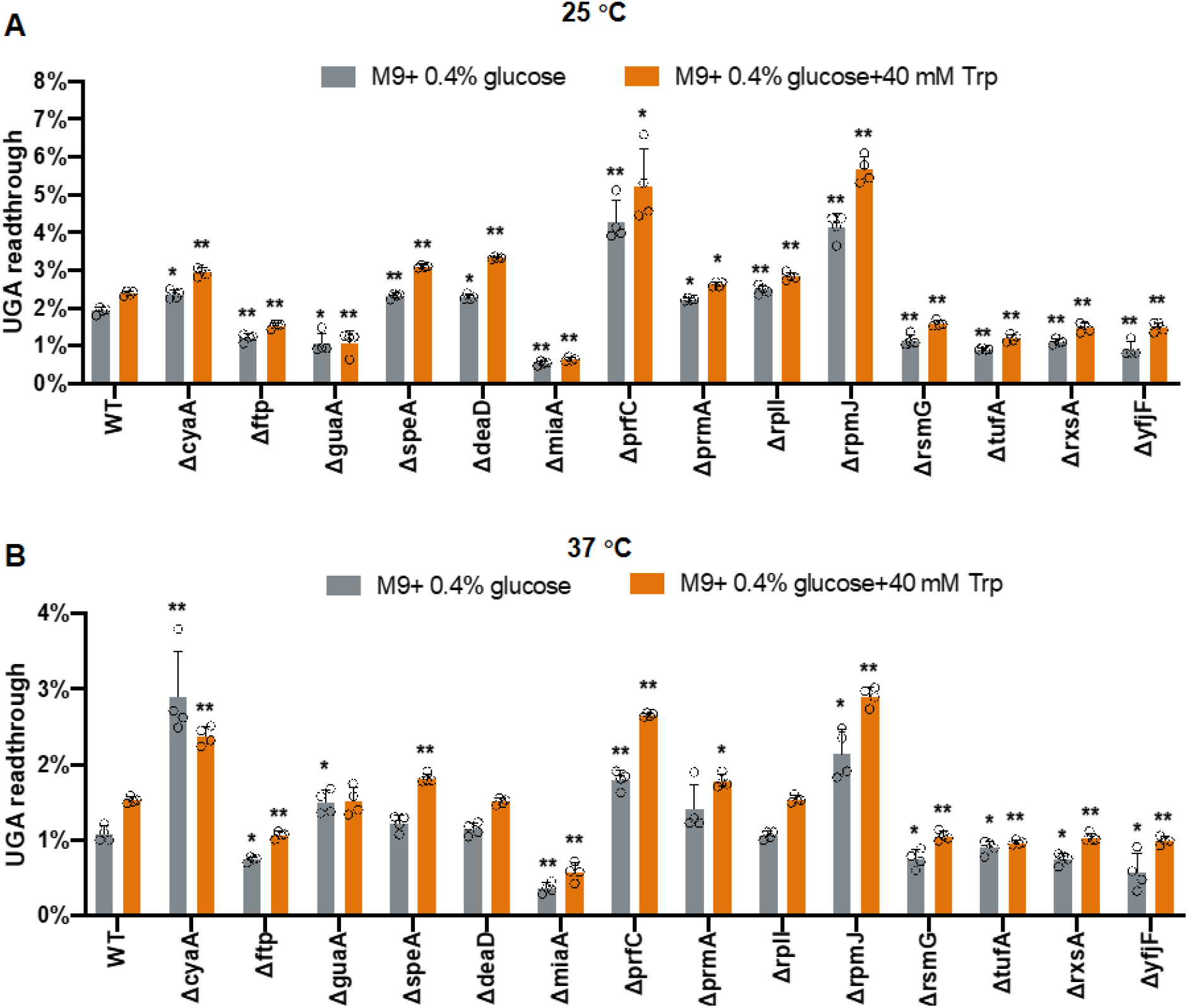
UGA readthrough of *E. coli* variants in minimal media. Overnight cultures of *E. coli* with pZS-Ptet-m-TGA-y and pZS-Ptet-m-y were diluted 1:50 in M9 media and incubated at 25 °C **(A)** or 37 °C **(B)** for 16 hours. The UGA readthrough levels were determined as in Figure 2. Error bars indicate standard deviations. P values were calculated using the unpaired t-test comparing the mutants with the WT. * P ≤ 0.05, ** P ≤ 0.0001.

## DISCUSSION

Early studies of streptomycin-resistant *E. coli* mutants have led to the discovery of the critical roles that ribosomal genes *rpsD* (uS4) and *rpsL* (uS12) play in maintaining translational fidelity (52,53). Random mutagenesis of *rpsL* and rRNA genes has later been performed to identify mutations that affect ribosomal fidelity (24,54,55). More recently, suppression of stop codons has attracted tremendous interest in the treatment of genetic diseases caused by nonsense mutations (56–61). Several studies have used luminescence-based assays to screen for small compounds that promote the readthrough of stop codons (61,62). In this work, we have used an economical and convenient dual-fluorescence reporter assay to perform systematic and genome-wide screening for genes that affect translational fidelity. Compared with transposon-based selection, screening of a defined knockout library allows us to identify genes that either increase or decrease translational errors. Our screening has revealed a total of 30 nonessential genes in *E. coli* that affect UGA readthrough and most of them have not been previously shown to regulate translational fidelity.

Among the identified genes are a new class involved in metabolism (*crp, cyaA, ftp, guaA, purA, speA*, and *ubiEFH*). Further experiments suggest that CyaA enhances the biogenesis of tRNAs and rRNAs, likely through transcriptional regulation by CRP/cAMP (Figures 2–7). GuaA also increases the levels of tRNAs, in addition to its role in GTP synthesis which affects the activity of RF3 (Figures 2–7). SpeA contributes to polyamine synthesis and the increase of UGA readthrough in Δ*speA* is abolished in the presence of spermidine (Figure 6), supporting the role of polyamines in maintaining translational fidelity. It remains unclear why other genes involved in polyamine synthesis have little effect on UGA readthrough. The roles of other identified genes in stop-codon readthrough remain to be elucidated in future studies. RpmJ is a nonessential ribosomal protein that maintains the rRNA structure (63). Deleting *rpmJ* increases stop-codon readthrough and amino acid misincorporation (Figure 3). It’s possible that the ribosome lacking RpmJ contains a more relaxed decoding center. Another interesting group of the identified genes is the *rsx* operon, which is involved in redox regulation (64). Oxidative stress has pleiotropic effects on translational fidelity (16,28,36). It remains to be determined how oxidative stress affects ribosomal fidelity.

Increasing evidence suggests that translational fidelity is tightly coupled with cellular metabolism. Competition between tRNAs affects the rates of amino acid misincorporation (8,65), and fluctuations in amino acid concentrations are predicted to affect the fidelity of aa-tRNA synthesis (4). Stop codon readthrough is affected by pH (14,66,67), sugar metabolism (14), as well as concentrations of amino acids and tRNAs (Figures S7-S8). Conversely, single cells with higher UGA readthrough appear to grow better in minimal media but are more sensitive to stress (12,14). Future work is needed to elucidate how regulation of translational fidelity affects cellular metabolism.

## MATERIALS AND METHODS

### Bacterial strains, plasmids, growth conditions, and reagents

All strains and plasmids used in this study are listed in Supplementary Table S2. The oligonucleotides used for gene disruption and plasmid construction are listed in Table S3. Gene knockout was performed as described (68). Unless otherwise noted, all bacteria used in this study were cultured in regular Lennox broth consisting of 10 g/L tryptone, 5 g/L sodium chloride, and 5 g/L yeast extract at 25 °C or 37 °C. The composition of M9 medium was 47.8 mM Na_2_HPO_4_, 22.0 mM KH_2_PO_4_, 18.7 mM (NH4)_2_SO_4_, 8.6 mM NaCl, 2.0 mM MgSO_4_, 0.1 mM CaCl_2_, and 0.40 % glucose with or without supplementation of 40 mM tryptophan. The concentrations of antibiotics used were as follows: 100 μg/mL ampicillin (Amp), 25 μg/mL chloramphenicol (Chl), and 50 μg/mL kanamycin (Kan). 10 mM arabinose was added for the induction of the lambda Red recombinase on pKD46.

### Plasmid transformation

For the initial screening, plasmids were transformed into the KEIO collection in 96-well plates. Cells were grown at 37 °C in 100 μL LB per well to mid-log phase and centrifuged at 4000 g for 10 min. Each pellet was resuspended in 20 μL fresh transformation buffer (LB broth with 10% polyethylene glycol (PEG3350), 5% dimethyl sulfoxide (DMSO), 10 mM MgSO_4_, and 10 mM MgCl_2_). 20 μL KCM buffer (0.1 M KCl, 0.03 M CaCl_2_ and 0.05 M MgCl_2_) with ~10 ng plasmid was added to the competent cells, followed by 30 min of incubation on ice. The cell/DNA mixture was placed at room temperature for 10 min before the addition of 40 μL LB pre-warmed at 37 °C. The cells were then incubated at 37°C with shaking for 1 h. 80 μL LB with 200 μg/mL Amp was added to each well to select the transformants. After overnight incubation at 37°C, the cultures were diluted 1:50 in LB with 100 μg/mL Amp and incubated at 37°C with shaking to saturation.

### Fluorescence-based stop codon readthrough assay

Stop codon readthrough rates were determined as described (12). Briefly, overnight cultures of *E. coli* were diluted 1:50 in LB or M9 and incubated at 25 °C or 37 °C using 96-well black side plates with a clear bottom (Corning). After 16 h, the fluorescence intensity of mCherry and YFP was determined in a microplate reader (Synergy HTX, BioTek). The level of stop-codon readthrough was calculated by determining the YFP/mCherry ratio of pZS-Ptet-m-TGA-y relative to the YFP/mCherry ratio of the control pZS-Ptet-m-y, and pZS-Ptet-lacZ was used as for background subtraction. When necessary, 5 mM spermidine, 4 mM cyclic-AMP, 1 mM guanosine, 1% glucose, or 2 μg/mL Chl was added into the culture.

### Dual luciferase assay

Cells grown at 37°C to mid-log phase in LB supplemented with 100 μg/mL ampicillin were pelleted, washed once with phosphate-buffered saline (PBS), and lysed in 50 μL passive lysis buffer (PLB). After incubation at room temperature for 15 min, the samples were flash frozen in liquid nitrogen, and then thawed on ice. Firefly and Renilla luciferase activities were determined using the Dual-Luciferase Reporter Assay System (Promega) according to the manufacturer’s instruction and the error frequencies were determined as previously described (8).

### Single-cell UGA readthrough using fluorescence microscopy

Overnight cultures of *E. coli* carrying the pZS-Ptet-mCherry-TGA-y plasmid were diluted 1:50 in LB and grown aerobically at 25 °C for 16 h with shaking. 3 μL of the resulting cultures were placed on a 1.5% agarose LB pad on a single slide. Fluorescence images were obtained using BZ-X800 fluorescence microscope (Keyence) with a 100x oil lens.

### Measurements of promoter activity with a platereader

Overnight cultures were diluted 1:50 and grown in LB at 25 °C or 37 °C with vigorous shaking. After 16 h, the fluorescence was measured using the microplate reader 3 (Synergy HTX, BioTek) as follows: mCherry (excitation wavelength at 575 nm and emission wavelength at 620 nm), and eCFP (excitation wavelength at 420 nm and emission wavelength at 475 nm). The gain was set manually to 40. The promoter activity was calculated as the ratio of eCFP over mCherry.

### RNA/protein ratio assay

The measurement of RNA/protein ratios was adapted from You et al (69). Briefly, cells were grown at 25 °C or 37 °C to the mid-log phase in LB with shaking. 1.5 mL culture was collected by centrifugation and the cell pellet was fast frozen in liquid nitrogen. For the total RNA quantification, the pellet was first washed twice with 0.7 M HClO_4_ and digested with 0.3 M KOH for 1 h at 37 °C with occasional mixing. The cell extract was further neutralized with 0.1 ml of 3 M HClO_4_, and the supernatant was collected. The precipitate was re-extracted twice with 0.55 mL of 0.5 M HClO_4_ and a final volume of 1.5-mL supernatant was combined. The total RNA concentration was determined in a microplate reader Take 3 (Synergy HTX, BioTek). For quantification of total proteins, the thawed cell pellet was digested with 100 μL of 3 M NaOH and heated at 98 °C for 5 min before cooling down to room temperature. The Biuret reaction was carried out by adding 100 μL of 1.6% CuSO_4_ to the above cell mixture with thorough mixing and incubating at room temperature for 5 min. The reaction mixture was centrifuged at 13,000 g for 1 min and the supernatant was measured for its absorbance at 555 nm. A similar experimental process was applied to a series of bovine serum albumin (BSA) standards to obtain a standard curve.

### Northern blot analysis

The total RNA was isolated from 1 mL mid-log phase cultures using TRIzol® Reagent (Invitrogen). The resulting lysate was phase separated with 200 μL chloroform and the total RNA was precipitated with the same volume of ice-cold isopropanol. The pellet was washed twice with 75% ethanol and resuspended in 30-50 μL of nuclease-free water by incubating at room temperature for 20 min. 3 μg total RNA was separated by electrophoresis on a vertical 8% (w/v) polyacrylamide gel with 8 M urea in Tris-Borate-EDTA (TBE) buffer at 120 V for 3 h. The samples were then transferred to the Zeta-Probe nylon membrane (Bio-Rad), cross-linked by UV light exposure, and hybridized with oligonucleotide probes labeled with biotin at the 5’ end (see Table S3) at 42 °C. The tRNA^Trp^ signal was normalized to the signal of the loading control SsrA RNA by using ImageJ (The National Institutes of Health).

### Determination of growth rate

*E. coli* cells were diluted 1:100 into fresh LB and incubated at 25 °C or 37 °C for 16 h with shaking. Cell growth was automatically monitored every 20 min by measuring the optical density at 600 nm (A600) using a microplate reader (Synergy HTX, BioTek). The growth rates were obtained from the exponential growth phase.

### Intracellular ATP detection

Cells were grown at 37 °C in LB to mid-log phase and were normalized by A600. 1 mL culture aliquot was washed once with ice-cold PBS and resuspended in 300 μL of the same buffer. The ATP level was measured using an ATP Determination Kit (Invitrogen) according to the manufacturer’s instructions. Blank PBS was used for background subtraction.

### Flow cytometry

Cells harboring pZS-Ptet-eCFP-TGA-y plasmid were grown in fresh LB aerobically for 16 h at 25 °C with shaking, diluted in PBS with 25 μg/mL Chl, and kept at 4 °C before flow cytometry on BD FACSCanto II flow cytometer. 30000 gated events were collected for each sample and FlowJo software was used for further data analysis.

### Polysome profiling

Polysome profiling was performed essentially as described (70). *E. coli* cells were grown in 100 mL LB to mid-log phase. Translation was then inhibited using 100 μg/mL Chl and quickly placing the cultures on ice for 10 minutes. Cells were then collected and washed twice with lysis buffer (20 mM Tris HCl pH 8, 140 mM KCl, 40 mM MgCl_2_, 0.5 mM dithiothreitol, 100 μg/mL Chl, and 1 mg/mL heparin). Cells were then lysed using glass beads. Samples were standardized on A260 readings. 150 A260 units were loaded onto 10%-50% sucrose gradients prepared as described (71). Gradients were centrifuged using a SW40-TI rotor for 2.5 hours at 35,000 rpm at 4 °C. Polysome profiles were then analyzed by continuous reading at 254 nm.

### Statistical analyses

Experiments were performed using at least three biological replicates. In all cases, error bars represent the standard deviations (SD). Statistical differences were analyzed using the unpaired t-test. Differences were considered significant at a P value < 0.05.

## Supporting information

Supplemental Files Summary

## DATA AVAILABILITY

The data that support the findings of this study can be made available from the corresponding author upon reasonable request.

## SUPPLEMENTARY MATERIALS

Figures S1-S10. Tables S1-S3.

## ACKOWLEGMENTS

We would like to thank Dr. Philip J. Farabaugh (University of Maryland) for providing the dual-luciferase constructs pEK 4 and pEK7.

## AUTHOR CONTRIBUTIONS

Z.L, P.V., L.O., P.M., and J.L. designed, performed, and analyzed the experiments. Z.L. and J.L. wrote the manuscript. All authors proofread the manuscript.

## REFERENCES

1. Zaher, H.S. and Green, R. (2009) Fidelity at the molecular level: lessons from protein synthesis. Cell, 136, 746–762.

2. Ling, J., Reynolds, N. and Ibba, M. (2009) Aminoacyl-tRNA synthesis and translational quality control. Annu Rev Microbiol, 63, 61–78.

3. Rozov, A., Demeshkina, N., Westhof, E., Yusupov, M. and Yusupova, G. (2015) Structural insights into the translational infidelity mechanism. Nat Commun, 6, 7251.

4. Rubio Gomez, M.A. and Ibba, M. (2020) Aminoacyl-tRNA synthetases. RNA, 26, 910–936.

5. Kuzmishin Nagy, A.B., Bakhtina, M. and Musier-Forsyth, K. (2020) Trans-editing by aminoacyl-tRNA synthetase-like editing domains. Enzymes, 48, 69–115.

6. Martinis, S.A. and Boniecki, M.T. (2010) The balance between pre- and post-transfer editing in tRNA synthetases. FEBS Lett, 584, 455–459.

7. Rodnina, M.V. and Wintermeyer, W. (2001) Ribosome fidelity: tRNA discrimination, proofreading and induced fit. Trends Biochem Sci, 26, 124–130.

8. Kramer, E.B. and Farabaugh, P.J. (2007) The frequency of translational misreading errors in *E. coli* is largely determined by tRNA competition. RNA, 13, 87–96.

9. Mohler, K., Aerni, H.R., Gassaway, B., Ling, J., Ibba, M. and Rinehart, J. (2017) MS-READ: Quantitative measurement of amino acid incorporation. Biochim Biophys Acta, 1861, 3081–3088.

10. Mordret, E., Dahan, O., Asraf, O., Rak, R., Yehonadav, A., Barnabas, G.D., Cox, J., Geiger, T., Lindner, A.B. and Pilpel, Y. (2019) Systematic detection of amino acid substitutions in proteomes reveals mechanistic basis of ribosome errors and selection for translation fidelity. Mol Cell, 75, 427–441 e425.

11. Devaraj, A., Shoji, S., Holbrook, E.D. and Fredrick, K. (2009) A role for the 30S subunit E site in maintenance of the translational reading frame. RNA., 15, 255–265.

12. Fan, Y., Evans, C.R., Barber, K.W., Banerjee, K., Weiss, K.J., Margolin, W., Igoshin, O.A., Rinehart, J. and Ling, J. (2017) Heterogeneity of stop codon readthrough in single bacterial cells and implications for population fitness. Mol Cell, 67, 826–836.

13. Engelberg-Kulka, H. (1981) UGA suppression by normal tRNA^Trp^ in *Escherichia coli*: codon context effects. Nucleic Acids Res, 9, 983–991.

14. Zhang, H., Lyu, Z., Fan, Y., Evans, C.R., Barber, K.W., Banerjee, K., Igoshin, O.A., Rinehart, J. and Ling, J. (2020) Metabolic stress promotes stop-codon readthrough and phenotypic heterogeneity. Proc Natl Acad Sci USA, 117, 22167–22172.

15. Schwartz, M.H., Waldbauer, J.R., Zhang, L. and Pan, T. (2016) Global tRNA misacylation induced by anaerobiosis and antibiotic exposure broadly increases stress resistance in *Escherichia coli*. Nucleic Acids Res, 44, 10292–10303.

16. Steiner, R.E., Kyle, A.M. and Ibba, M. (2019) Oxidation of phenylalanyl-tRNA synthetase positively regulates translational quality control. Proc Natl Acad Sci USA, 116, 10058–10063.

17. Lant, J.T., Kiri, R., Duennwald, M.L. and O’Donoghue, P. (2021) Formation and persistence of polyglutamine aggregates in mistranslating cells. Nucleic Acids Res, 49, 11883–11899.

18. Schmidt, E. and Schimmel, P. (1994) Mutational isolation of a sieve for editing in a transfer RNA synthetase. Science, 264, 265–267.

19. Roy, H., Ling, J., Irnov, M. and Ibba, M. (2004) Post-transfer editing *in vitro* and *in vivo* by the beta subunit of phenylalanyl-tRNA synthetase. EMBO J, 23, 4639–4648.

20. Beuning, P.J. and Musier-Forsyth, K. (2000) Hydrolytic editing by a class II aminoacyl-tRNA synthetase. Proc Natl Acad Sci USA, 97, 8916–8920.

21. Dock-Bregeon, A., Sankaranarayanan, R., Romby, P., Caillet, J., Springer, M., Rees, B., Francklyn, C.S., Ehresmann, C. and Moras, D. (2000) Transfer RNA-mediated editing in threonyl-tRNA synthetase. The class II solution to the double discrimination problem. Cell, 103, 877–884.

22. Ling, J., O’Donoghue, P. and Söll, D. (2015) Genetic code flexibility in microorganisms: novel mechanisms and impact on physiology. Nat Rev Microbiol, 13, 707–721.

23. Lant, J.T., Berg, M.D., Heinemann, I.U., Brandl, C.J. and O’Donoghue, P. (2019) Pathways to disease from natural variations in human cytoplasmic tRNAs. J Biol Chem, 294, 5294–5308.

24. McClory, S.P., Leisring, J.M., Qin, D. and Fredrick, K. (2010) Missense suppressor mutations in 16S rRNA reveal the importance of helices h8 and h14 in aminoacyl-tRNA selection. RNA, 16, 1925–1934.

25. Zimmermann, R.A., Garvin, R.T. and Gorini, L. (1971) Alteration of a 30S ribosomal protein accompanying the ram mutation in *Escherichia coli*. Proc Natl Acad Sci USA, 68, 2263–2267.

26. Dinman, J.D. (2019) Translational recoding signals: Expanding the synthetic biology toolbox. J Biol Chem, 294, 7537–7545.

27. Choi, J., O’Loughlin, S., Atkins, J.F. and Puglisi, J.D. (2020) The energy landscape of −1 ribosomal frameshifting. Sci Adv, 6, eaax6969.

28. Ling, J. and Söll, D. (2010) Severe oxidative stress induces protein mistranslation through impairment of an aminoacyl-tRNA synthetase editing site. Proc Natl Acad Sci USA, 107, 4028–4033.

29. Ballesteros, M., Fredriksson, A., Henriksson, J. and Nystrom, T. (2001) Bacterial senescence: protein oxidation in non-proliferating cells is dictated by the accuracy of the ribosomes. EMBO J, 20, 5280–5289.

30. Mohler, K. and Ibba, M. (2017) Translational fidelity and mistranslation in the cellular response to stress. Nat Microbiol, 2, 17117.

31. Martinez-Miguel, V.E., Lujan, C., Espie-Caullet, T., Martinez-Martinez, D., Moore, S., Backes, C., Gonzalez, S., Galimov, E.R., Brown, A.E.X., Halic, M. et al. (2021) Increased fidelity of protein synthesis extends lifespan. Cell Metab, 33, 2288–2300 e2212.

32. Shcherbakov, D., Nigri, M., Akbergenov, R., Brilkova, M., Mantovani, M., Petit, P.I., Grimm, A., Karol, A.A., Teo, Y., Sanchon, A.C. et al. (2022) Premature aging in mice with error-prone protein synthesis. Sci Adv, 8, eabl9051.

33. Fan, Y., Wu, J., Ung, M.H., De Lay, N., Cheng, C. and Ling, J. (2015) Protein mistranslation protects bacteria against oxidative stress. Nucleic Acids Res, 43, 1740–1748.

34. Fan, Y., Thompson, L., Lyu, Z., Cameron, T.A., De Lay, N.R., Krachler, A.M. and Ling, J. (2019) Optimal translational fidelity is critical for *Salmonella* virulence and host interactions. Nucleic Acids Res, 47, 5356–5367.

35. Javid, B., Sorrentino, F., Toosky, M., Zheng, W., Pinkham, J.T., Jain, N., Pan, M., Deighan, P. and Rubin, E.J. (2014) Mycobacterial mistranslation is necessary and sufficient for rifampicin phenotypic resistance. Proc Natl Acad Sci USA, 111, 1132–1137.

36. Netzer, N., Goodenbour, J.M., David, A., Dittmar, K.A., Jones, R.B., Schneider, J.R., Boone, D., Eves, E.M., Rosner, M.R., Gibbs, J.S. et al. (2009) Innate immune and chemically triggered oxidative stress modifies translational fidelity. Nature, 462, 522–526.

37. Lee, J.Y., Kim, D.G., Kim, B.G., Yang, W.S., Hong, J., Kang, T., Oh, Y.S., Kim, K.R., Han, B.W., Hwang, B.J. et al. (2014) Promiscuous methionyl-tRNA synthetase mediates adaptive mistranslation to protect cells against oxidative stress. J Cell Sci, 127, 4234–4245.

38. Baba, T., Ara, T., Hasegawa, M., Takai, Y., Okumura, Y., Baba, M., Datsenko, K.A., Tomita, M., Wanner, B.L. and Mori, H. (2006) Construction of *Escherichia coli* K-12 in-frame, single-gene knockout mutants: the Keio collection. Mol Syst Biol, 2, 2006.0008.

39. Petrullo, L.A., Gallagher, P.J. and Elseviers, D. (1983) The role of 2-methylthio-N6-isopentenyladenosine in readthrough and suppression of nonsense codons in *Escherichia coli*. Mol Gen Genet, 190, 289–294.

40. Thompson, K.M. and Gottesman, S. (2014) The MiaA tRNA modification enzyme is necessary for robust RpoS expression in *Escherichia coli*. J Bacteriol, 196, 754–761.

41. Youngman, E.M., McDonald, M.E. and Green, R. (2008) Peptide release on the ribosome: mechanism and implications for translational control. Annu Rev Microbiol, 62, 353–373.

42. Herr, A.J., Nelson, C.C., Wills, N.M., Gesteland, R.F. and Atkins, J.F. (2001) Analysis of the roles of tRNA structure, ribosomal protein L9, and the bacteriophage T4 gene 60 bypassing signals during ribosome slippage on mRNA. J Mol Biol, 309, 1029–1048.

43. Seidman, J.S., Janssen, B.D. and Hayes, C.S. (2011) Alternative fates of paused ribosomes during translation termination. J Biol Chem, 286, 31105–31112.

44. Wong, S.Y., Javid, B., Addepalli, B., Piszczek, G., Strader, M.B., Limbach, P.A. and Barry, C.E., 3rd. (2013) Functional role of methylation of G518 of the 16S rRNA 530 loop by GidB in *Mycobacterium tuberculosis*. Antimicrob Agents Chemother, 57, 6311–6318.

45. Zheng, D., Constantinidou, C., Hobman, J.L. and Minchin, S.D. (2004) Identification of the CRP regulon using *in vitro* and *in vivo* transcriptional profiling. Nucleic Acids Res, 32, 5874–5893.

46. Petersen, C. (1999) Inhibition of cellular growth by increased guanine nucleotide pools. Characterization of an *Escherichia coli* mutant with a guanosine kinase that is insensitive to feedback inhibition by GTP. J Biol Chem, 274, 5348–5356.

47. Maas, W.K. (1972) Mapping of genes involved in the synthesis of spermidine in *Escherichia coli*. Mol Gen Genet, 119, 1–9.

48. Kitagawa, M., Ara, T., Arifuzzaman, M., Ioka-Nakamichi, T., Inamoto, E., Toyonaga, H. and Mori, H. (2005) Complete set of ORF clones of *Escherichia coli* ASKA library (a complete set of *E. coli* K-12 ORF archive): unique resources for biological research. DNA Res, 12, 291–299.

49. McMurry, L.M. and Algranati, I.D. (1986) Effect of polyamines on translation fidelity in vivo. Eur J Biochem, 155, 383–390.

50. Abraham, A.K., Olsnes, S. and Pihl, A. (1979) Fidelity of protein synthesis in vitro is increased in the presence of spermidine. FEBS Lett, 101, 93–96.

51. Dai, X., Zhu, M., Warren, M., Balakrishnan, R., Patsalo, V., Okano, H., Williamson, J.R., Fredrick, K., Wang, Y.P. and Hwa, T. (2016) Reduction of translating ribosomes enables *Escherichia coli* to maintain elongation rates during slow growth. Nat Microbiol, 2, 16231.

52. Davies, J., Gilbert, W. and Gorini, L. (1964) Streptomycin, suppression, and the code. Proc Natl Acad Sci USA, 51, 883–890.

53. Gorini, L. and Kataja, E. (1964) Phenotypic repair by streptomycin of defective genotypes in *E. coli*. Proc Natl Acad Sci USA, 51, 487–493.

54. Agarwal, D., Gregory, S.T. and O’Connor, M. (2011) Error-prone and error-restrictive mutations affecting ribosomal protein S12. J Mol Biol, 410, 1–9.

55. Qin, D., Abdi, N.M. and Fredrick, K. (2007) Characterization of 16S rRNA mutations that decrease the fidelity of translation initiation. RNA, 13, 2348–2355.

56. Ng, M.Y., Li, H., Ghelfi, M.D., Goldman, Y.E. and Cooperman, B.S. (2021) Ataluren and aminoglycosides stimulate read-through of nonsense codons by orthogonal mechanisms. Proc Natl Acad Sci USA, 118.

57. Roy, B., Friesen, W.J., Tomizawa, Y., Leszyk, J.D., Zhuo, J., Johnson, B., Dakka, J., Trotta, C.R., Xue, X., Mutyam, V. et al. (2016) Ataluren stimulates ribosomal selection of near-cognate tRNAs to promote nonsense suppression. Proc Natl Acad Sci USA, 113, 12508–12513.

58. Wang, J., Zhang, Y., Mendonca, C.A., Yukselen, O., Muneeruddin, K., Ren, L., Liang, J., Zhou, C., Xie, J., Li, J. et al. (2022) AAV-delivered suppressor tRNA overcomes a nonsense mutation in mice. Nature, 604, 343–348.

59. Beznoskova, P., Bidou, L., Namy, O. and Valasek, L.S. (2021) Increased expression of tryptophan and tyrosine tRNAs elevates stop codon readthrough of reporter systems in human cell lines. Nucleic Acids Res, 49, 5202–5215.

60. Albers, S., Beckert, B., Matthies, M.C., Mandava, C.S., Schuster, R., Seuring, C., Riedner, M., Sanyal, S., Torda, A.E., Wilson, D.N. et al. (2021) Repurposing tRNAs for nonsense suppression. Nat Commun, 12, 3850.

61. Welch, E.M., Barton, E.R., Zhuo, J., Tomizawa, Y., Friesen, W.J., Trifillis, P., Paushkin, S., Patel, M., Trotta, C.R., Hwang, S. et al. (2007) PTC124 targets genetic disorders caused by nonsense mutations. Nature, 447, 87–91.

62. Liang, F., Shang, H., Jordan, N.J., Wong, E., Mercadante, D., Saltz, J., Mahiou, J., Bihler, H.J. and Mense, M. (2017) High-throughput screening for readthrough modulators of CFTR PTC mutations. SLAS Technol, 22, 315–324.

63. Maeder, C. and Draper, D.E. (2005) A small protein unique to bacteria organizes rRNA tertiary structure over an extensive region of the 50 S ribosomal subunit. J Mol Biol, 354, 436–446.

64. Koo, M.S., Lee, J.H., Rah, S.Y., Yeo, W.S., Lee, J.W., Lee, K.L., Koh, Y.S., Kang, S.O. and Roe, J.H. (2003) A reducing system of the superoxide sensor SoxR in *Escherichia coli*. EMBO J, 22, 2614–2622.

65. Kramer, E.B., Vallabhaneni, H., Mayer, L.M. and Farabaugh, P.J. (2010) A comprehensive analysis of translational missense errors in the yeast *Saccharomyces cerevisiae*. RNA, 16, 1797–1808.

66. Kuhlenkoetter, S., Wintermeyer, W. and Rodnina, M.V. (2011) Different substrate-dependent transition states in the active site of the ribosome. Nature, 476, 351–354.

67. Shaw, J.J., Trobro, S., He, S.L., Aqvist, J. and Green, R. (2012) A Role for the 2’ OH of peptidyl-tRNA substrate in peptide release on the ribosome revealed through RF-mediated rescue. Chem Biol, 19, 983–993.

68. Datsenko, K.A. and Wanner, B.L. (2000) One-step inactivation of chromosomal genes in *Escherichia coli* K-12 using PCR products. Proc Natl Acad Sci USA, 97, 6640–6645.

69. You, C., Okano, H., Hui, S., Zhang, Z., Kim, M., Gunderson, C.W., Wang, Y.P., Lenz, P., Yan, D. and Hwa, T. (2013) Coordination of bacterial proteome with metabolism by cyclic AMP signalling. Nature, 500, 301–306.

70. Nguyen, H.L., Duviau, M.P., Laguerre, S., Nouaille, S., Cocaign-Bousquet, M. and Girbal, L. (2022) Synergistic regulation of transcription and translation in *Escherichia coli* revealed by codirectional increases in mRNA concentration and translation efficiency. Microbiol Spectr, 10, e0204121.

71. Qin, D. and Fredrick, K. (2013) Analysis of polysomes from bacteria. Methods Enzymol, 530, 159–172.

