## Supplemental Files Summary for "Genome-wide screening reveals metabolic regulation of translational fidelity"

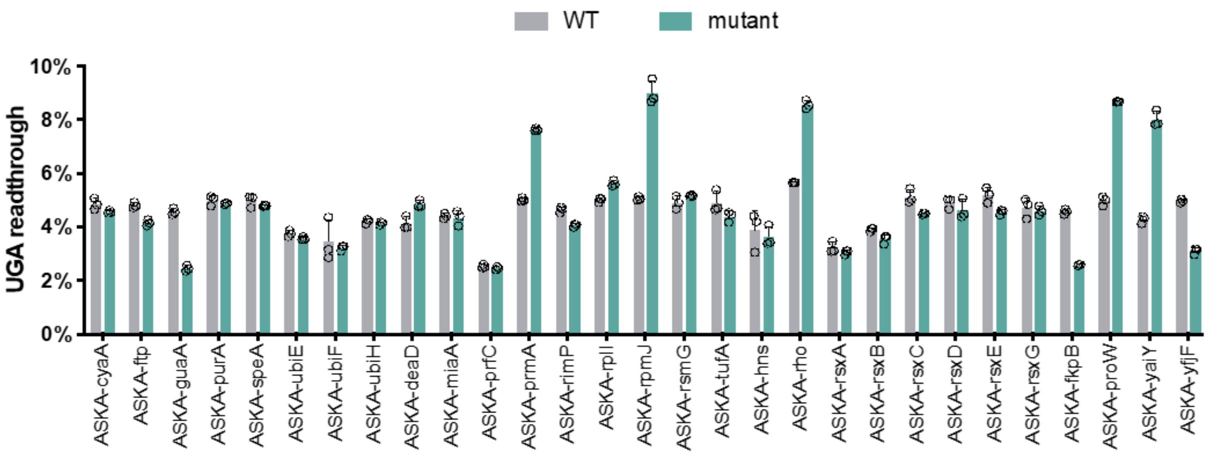


**Figure S1. UGA readthrough of *E. coli* variants complemented with overexpression plasmids.** Complementary plasmids from ASKA *E. coli* ORF library were transformed into *E. coli* BW25113 mutant strains with pZS-Ptet-m-TGA-y and pZS-Ptet-m-y. The UGA readthrough levels were quantified as in Figure 2. Error bars indicate standard deviations.

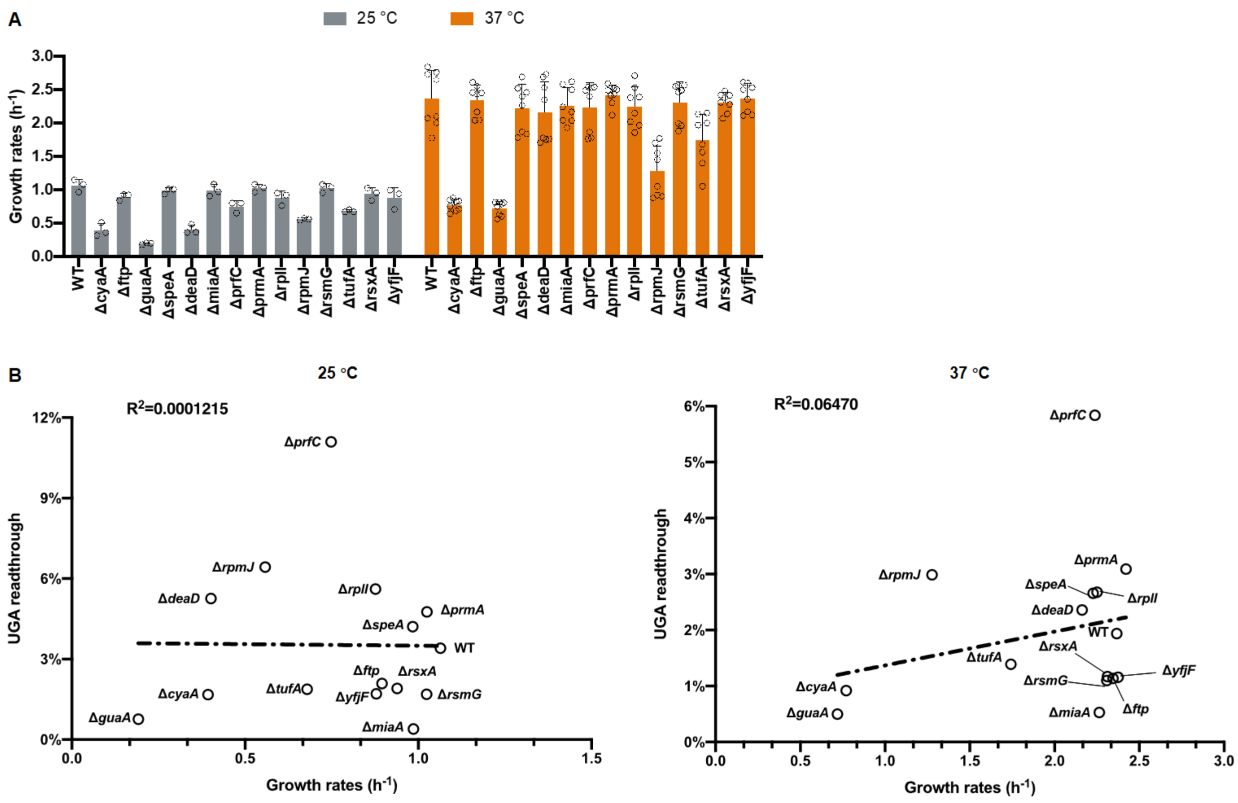


**Figure S2. Growth rates of MG1655-derived variants. (A)** Growth rates of *E. coli* variants in LB. **(B)** Growth rates are not correlated with UGA readthrough levels. Error bars indicate standard deviations.

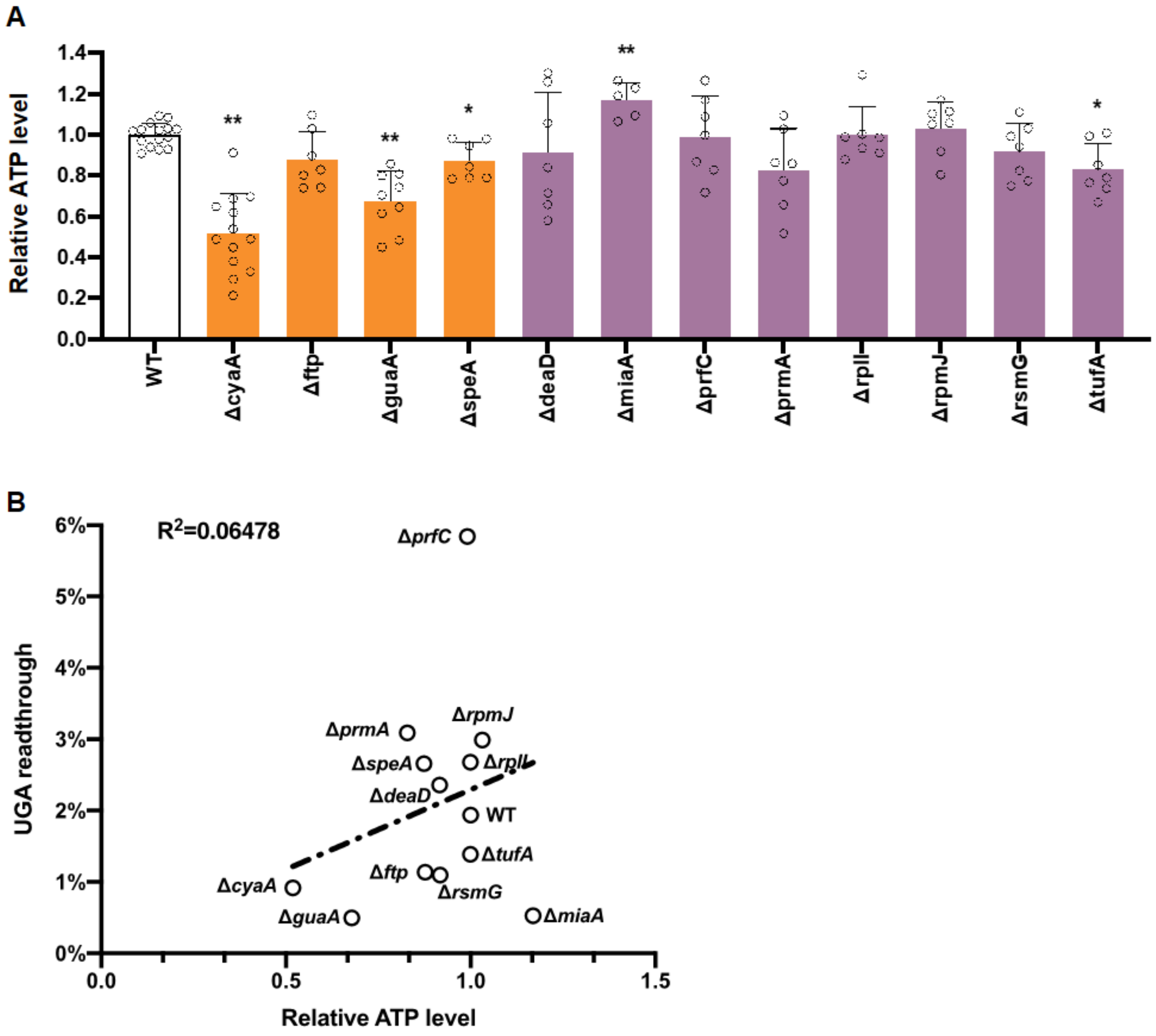


**Figure S3. Intracellular ATP levels. (A)** Relative ATP cells in cells grown at 37°C in LB to mid-log phase. **(B)** The UGA readthrough level does not appear to correlate with the ATP level. Error bars indicate standard deviations. P values were calculated using the unpaired t-test comparing the mutants with the WT. * $P \leqslant 0.05$, ** $P \leqslant 0.0001$.

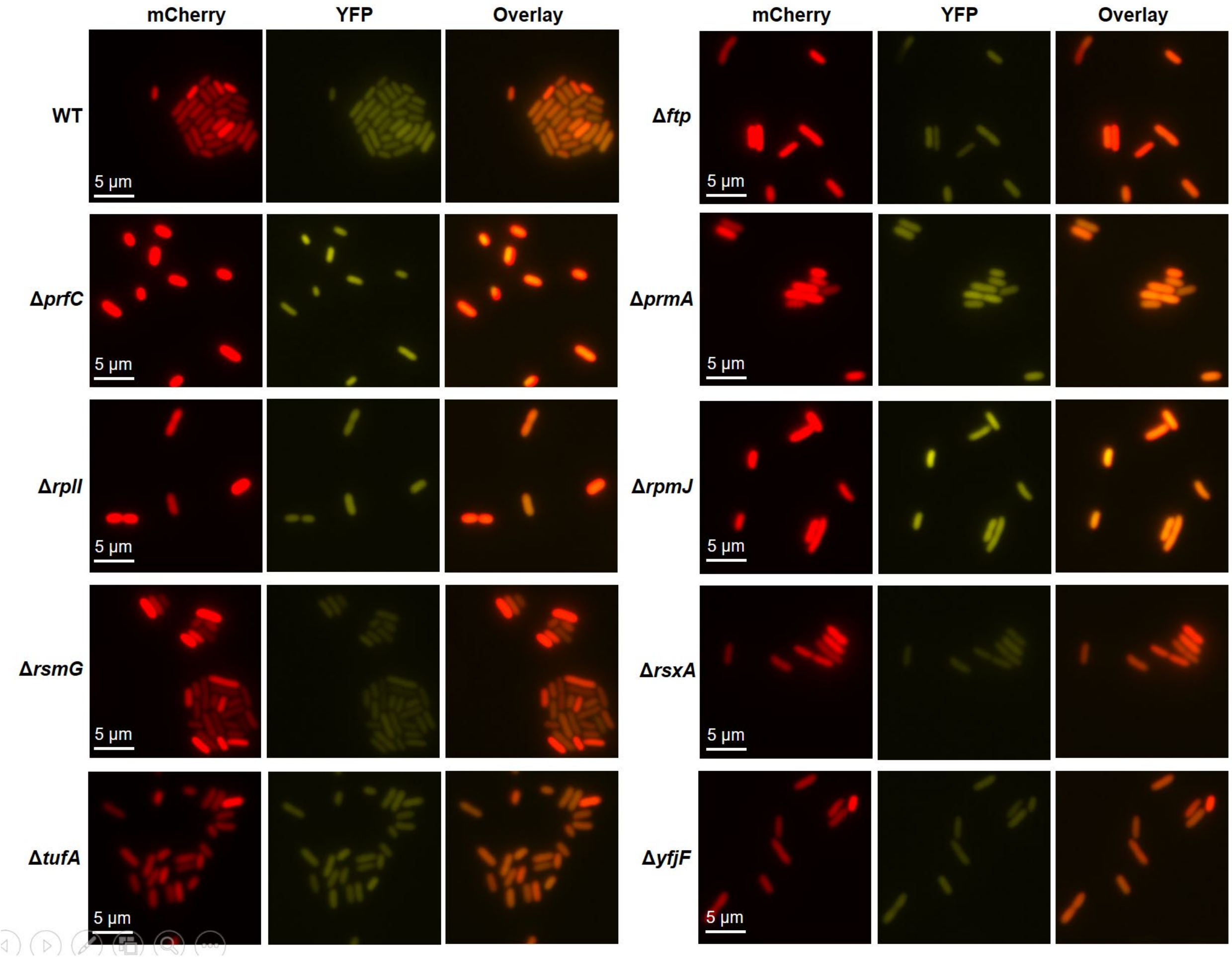


**Figure S4. UGA readthrough in single cells revealed by fluorescence microscopy**. The growth and imaging conditions were the same as in Figure 4.

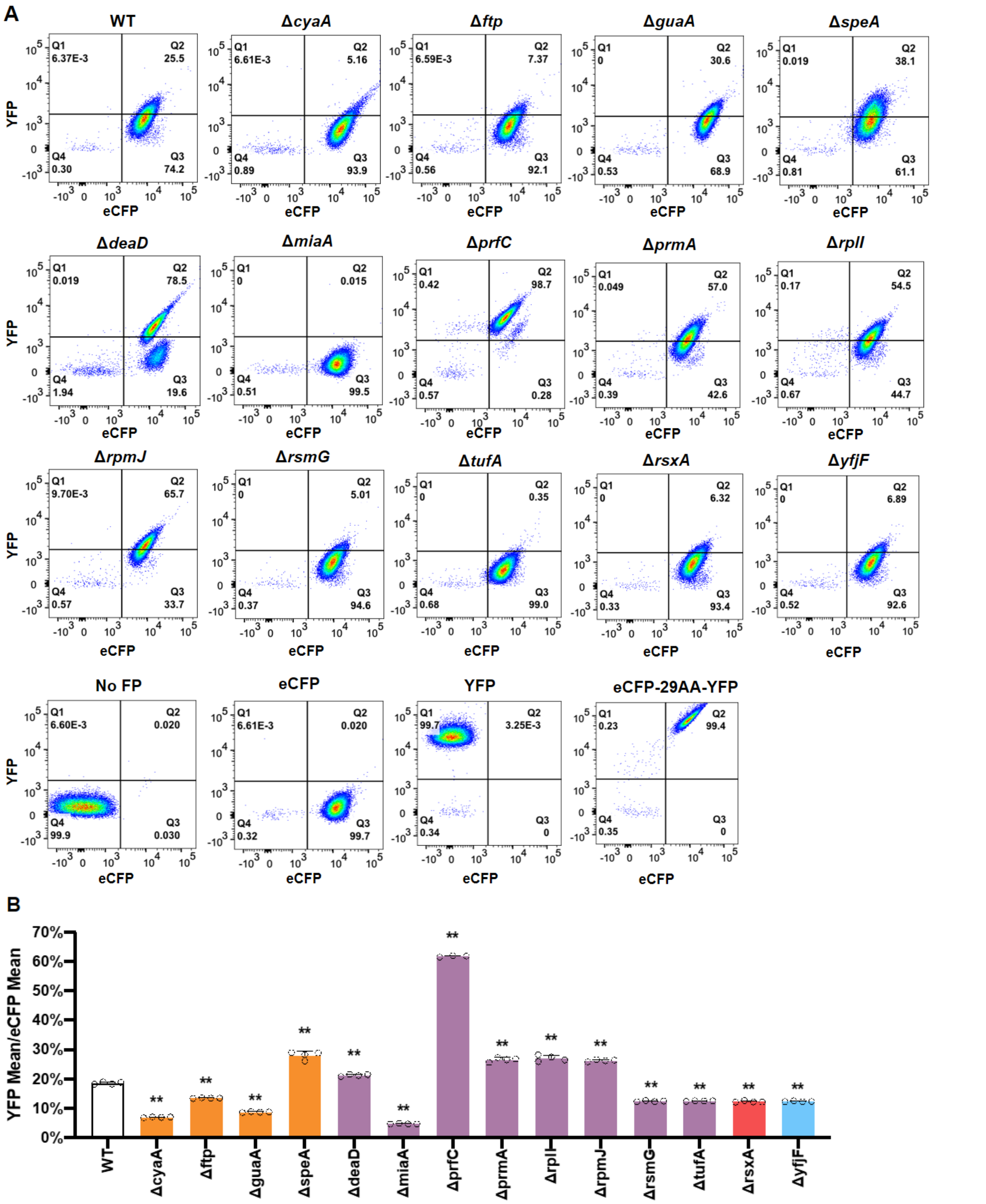


**Figure S5. UGA readthrough determined by flow cytometry.** Overnight cultures of *E. coli* with pZS-Ptet-eCFP-TGA-y were grown in LB Amp for 16 h at 25 °C. **(A)** Representative plots. **(B)** Mean YFP intensity divided by mean eCFP intensity for each strain. Error bars indicate standard deviations. P values were calculated using the unpaired t-test comparing the mutants with the WT. ** $P \leqslant 0.0001$.

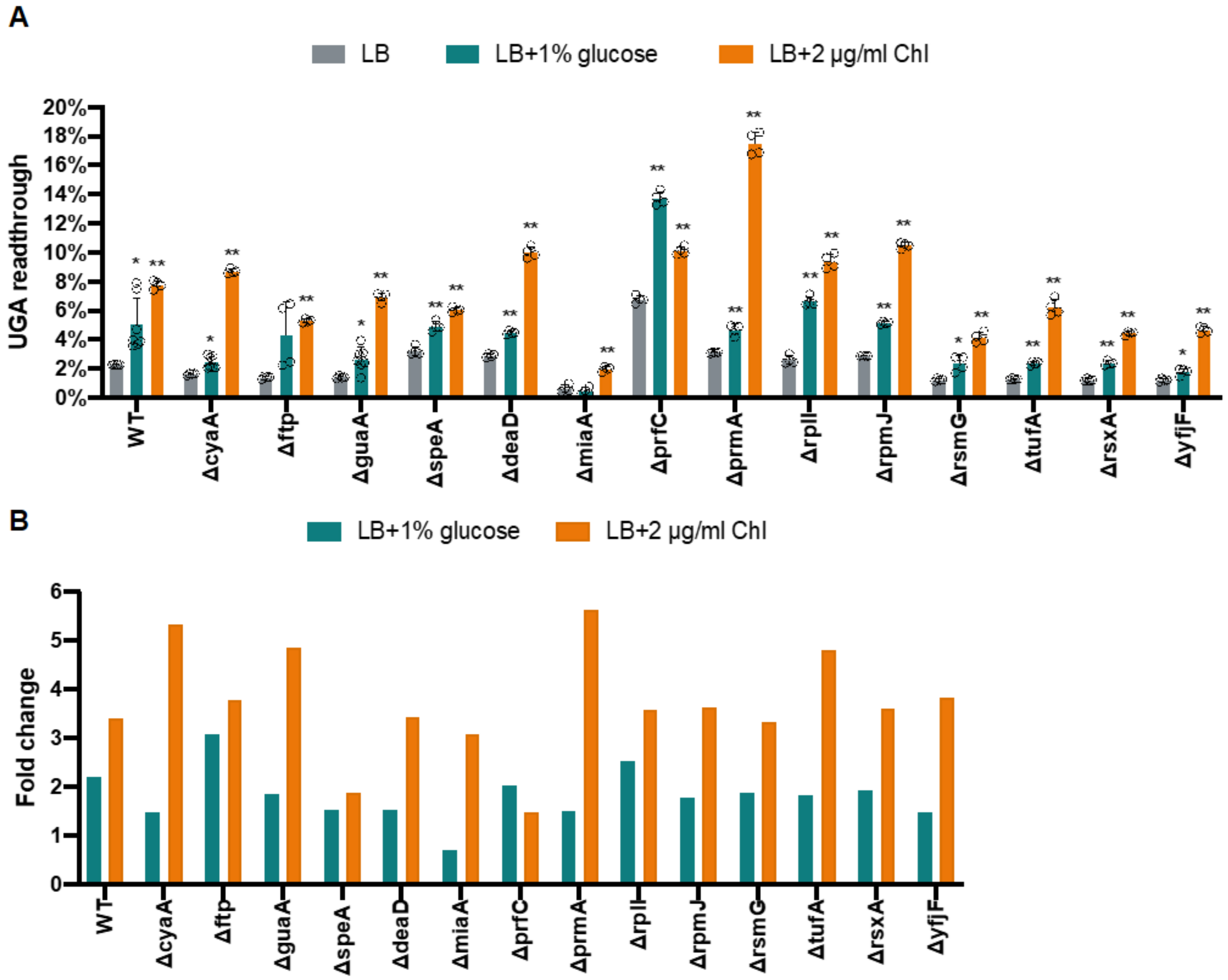


**Figure S6. UGA readthrough with glucose or chloramphenicol. (A)** Overnight cultures of *E. coli* with pZS-Ptet-m-TGA-y were diluted 1:50 in fresh LB Amp supplemented with 1% glucose or 2 μg/mL Chl and grown at 37 °C for 16 h. The UGA readthrough levels were determined as in Figure 2. **(B)** Fold change of UGA readthrough compared with that in LB. Error bars indicate standard deviations. P values were calculated using the unpaired t-test comparing the mutants with the WT grown in the same medium. * P ⩽ 0.05, ** P ⩽ 0.0001.

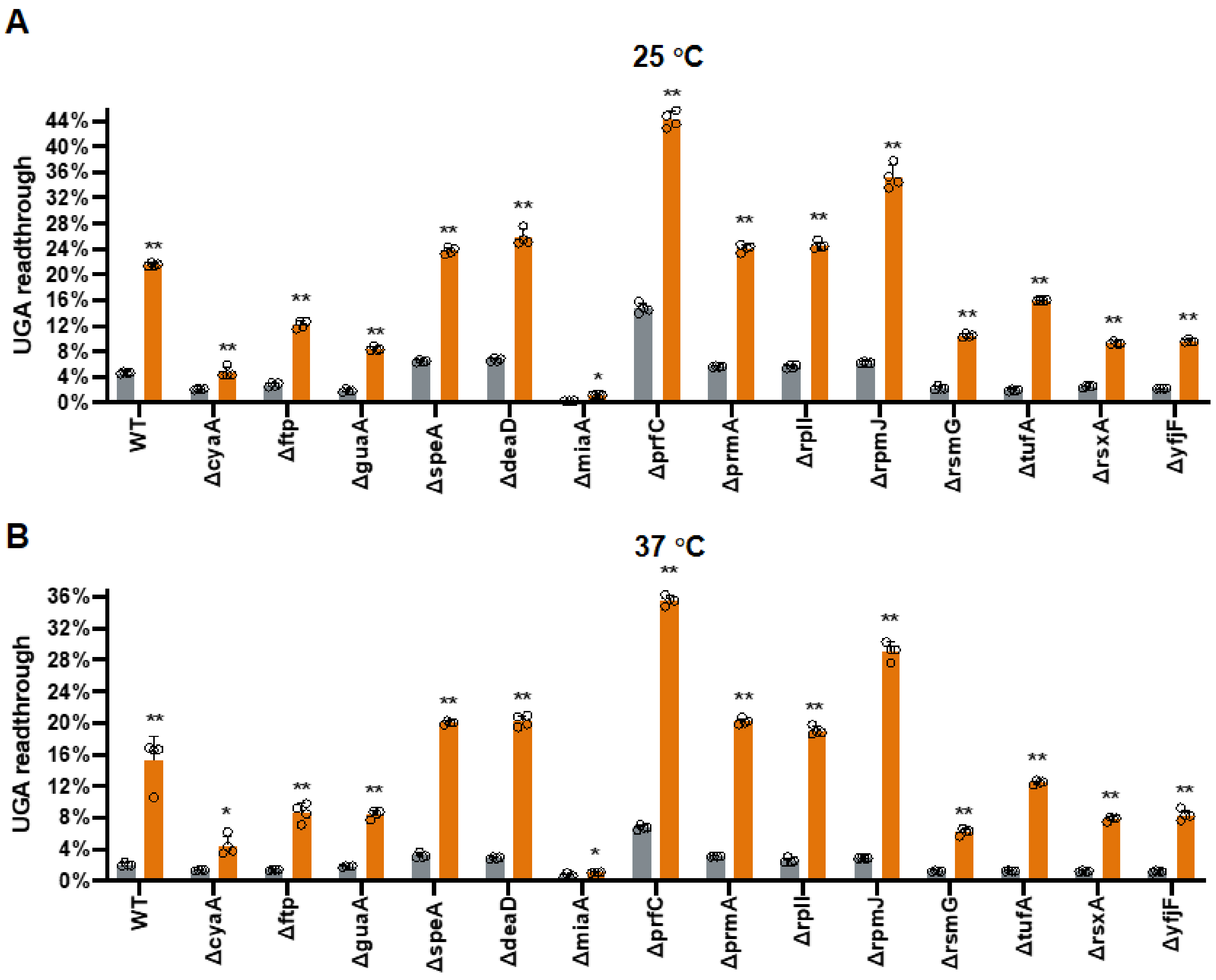


**Figure S7. Overexpressing tRNA$^{Trp}$ increases UGA readthrough.** The plasmid pKT tRNA$^{Trp}$ was transformed into *E. coli* variants. The growth condition at **(A)** 25 and **(B)** 37 °C and calculation of UGA readthrough were the same as in Figure 2. Grey, strains carrying empty pKT plasmid; orange, strains with pKT tRNA$^{Trp}$. Error bars indicate standard deviations. P values were calculated using the unpaired t-test comparing the mutants with the WT. * $P \leqslant 0.05$, ** $P \leqslant 0.0001$.

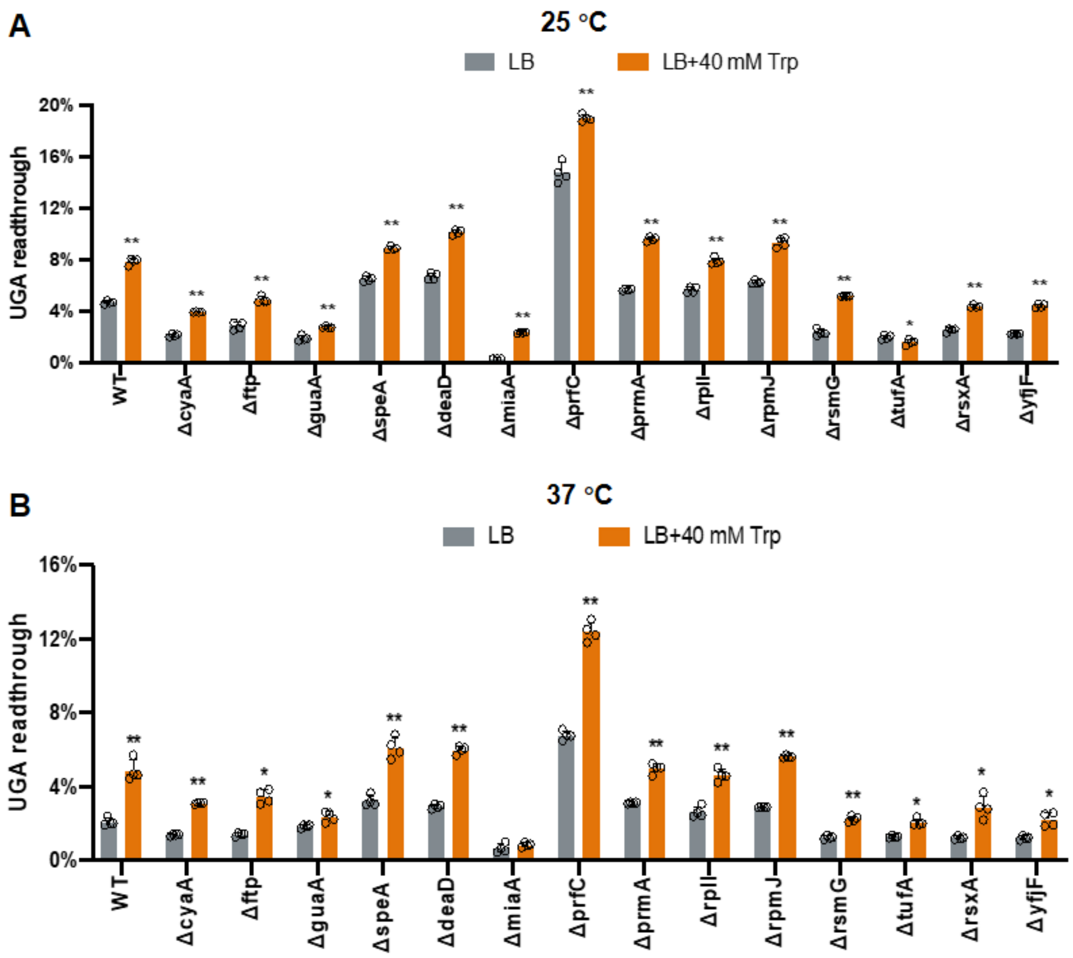


**Figure S8. Addition of Trp increases UGA readthrough.** The UGA readthrough levels were determined as in Figure 2 followed by growing cells at 25 °C **(A)** or 37 °C **(B)** for 16 h. Error bars indicate standard deviations. P values were calculated using the unpaired t-test comparing the mutants with the WT in the same medium. * $P \leqslant 0.05$, ** $P \leqslant 0.0001$.

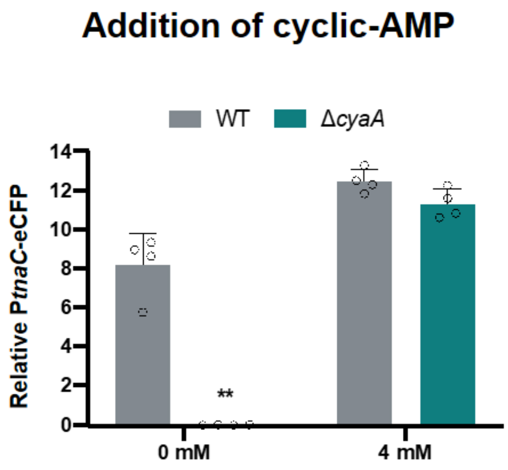


**Figure S9. Addition of cAMP restores activity of CRP/cAMP regulated promoter in Δ*cyaA*.** Overnight cultures harboring pZS-m-TGA-y-P*tnaC*-eCFP were diluted 1:50 in LB Amp with or without 4 mM cAMP and incubated at 25 °C for 16 h. The promoter activity was calculated as the ratio of eCFP over mCherry. The growth condition and calculation of UGA readthrough were the same as in Figure 2. Error bars indicate standard deviations. P values were calculated using the unpaired t-test comparing the mutants with the WT. ** P ⩽ 0.0001.

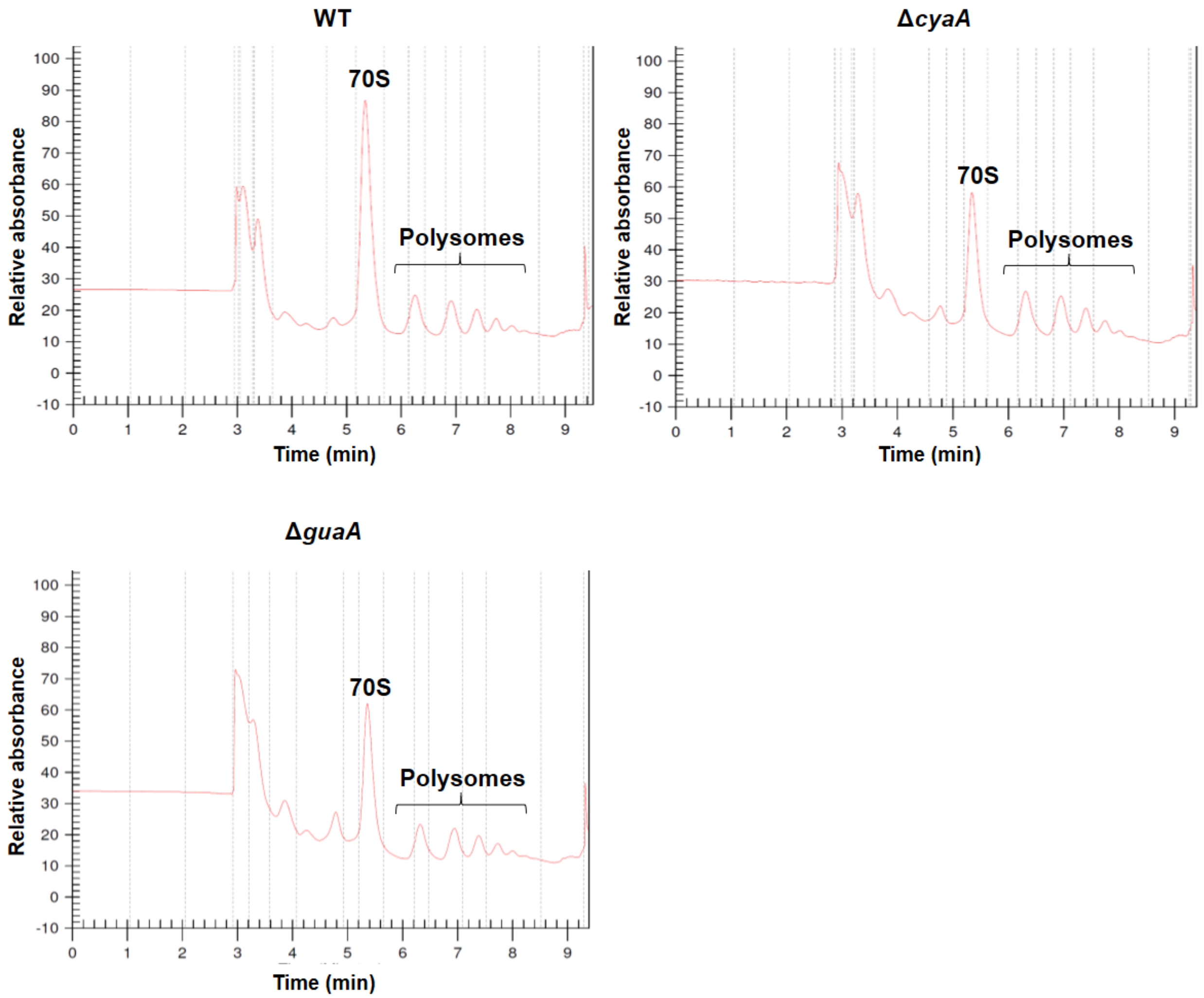


**Figure S10. Polysome profiling of WT, Δ*cyaA*, and Δ*guaA*.** The growth condition of the strains is described in MATERIALS AND METHODS.

**Table S1. List of identified genes from screening**

| GENE | DESCRIPTION |
|---|---|
| **Metabolism** | |
| *crp* | DNA-binding transcriptional dual regulator |
| *cyaA* | Adenylate cyclase |
| *guaA* | GMP synthetase |
| *purA* | Adenylosuccinate synthetase |
| *speA* | Biosynthetic arginine decarboxylase |
| *ftp* | FAD:protein FMN transferase |
| *ubiE* | Ubiquinone/menaquinone biosynthesis C-methyltransferase |
| *ubiF* | 3-demethoxyubiquinol 3-hydroxylase |
| *ubiH* | 2-octaprenyl-6-methoxyphenol 4-hydroxylase |
| **Translation** | |
| *deaD* | ATP-dependent RNA helicase |
| *miaA* | tRNA dimethylallyltransferase |
| *prfC* | Peptide chain release factor RF3 |
| *prmA* | Ribosomal protein L11 methyltransferase |
| *rimP* | Ribosome maturation factor |
| *rplI* | 50S ribosomal subunit protein L9 |
| *rpmJ* | 50S ribosomal subunit protein L36 |
| *rsmG* | 16S rRNA $m^7$G527 methyltransferase |
| *tufA* | Translation elongation factor Tu 1 |
| **Transcription** | |
| *hns* | DNA-binding transcriptional dual regulator |
| *rho* | Transcription termination factor |

| | |
|---|---|
| **Redox** | |
| *rsxABCDEFG* | SoxR [2Fe-2S] reducing system protein |
| **Other** | |
| *proW* | Glycine betaine ABC transporter membrane subunit |
| *fkpB* | Peptidyl-prolyl cis-trans isomerase |
| *yaiY* | DUF2755 domain-containing inner membrane protein |
| *yfjF* | Putative component of the Rsx system |

**Table S2. Strains and plasmids**

| Strains | Description | Source |
|---|---|---|
| Keio collection | *E. coli* K-12 BW25113 (F-, $\lambda^-$, rph-1) | (1) |
| MG1655 | *E. coli* K-12 MG1655 (F-, $\lambda^-$, rph-1) | Lab collection |
| Δ*cyaA* | MG Δ*cyaA*::FRT | This study |
| Δ*ftp* | MG Δ*ftp*::FRT | This study |
| Δ*guaA* | MG Δ*guaA*::FRT | This study |
| Δ*speA* | MG Δ*speA*::FRT | This study |
| Δ*deaD* | MG Δ*deaD*::FRT | This study |
| Δ*miaA* | MG Δ*miaA*::FRT | This study |
| Δ*prfC* | MG Δ*prfC*::FRT | This study |
| Δ*prmA* | MG Δ*prmA*::FRT | This study |
| Δ*rplI* | MG Δ*rplI*::FRT | This study |
| Δ*rpmJ* | MG Δ*rpmJ*::FRT | This study |
| Δ*rsmG* | M<br>G [illegible] | This study |
| Δ*tufA* | MG Δ*tufA*::FRT | This study |
| Δ*rsxA* | MG Δ*rsxA*::FRT | This study |
| Δ*fkpB* | MG Δ*fkpB*::FRT | This study |
| Δ*proW* | MG Δ*proW*::FRT | This study |
| Δ*yaiY* | MG Δ*yaiY*::FRT | This study |
| Δ*yfjF* | MG Δ*yfjF*::FRT | This study |
| Δ*cyaA*Δ*prfC* | G Δ*cyaA*::FRT Δ*prfC*::*cat* | This study |
| Δ*ftp* Δ*prfC* | MG Δ*ftp*::FRT Δ*prfC*::*cat* | This study |

| | | |
|---|---|---|
| Δ*guaA* Δ*prfC* | MG [illegible] | This study |
| Δ*speA* Δ*prfC* | MG Δ*speA*::FRT Δ*prfC*::*cat* | This study |
| Δ*deaD* Δ*prfC* | MG Δ*deaD*::FRT Δ*prfC*::*cat* | This study |
| Δ*miaA* Δ*prfC* | MG Δ*prfC*::FRT Δ*miaA*::*cat* | This study |
| Δ*prmA* Δ*prfC* | MG Δ*prmA*::FRT Δ*prfC*::*cat* | This study |
| Δ*rplI* Δ*prfC* | MG Δ*prfC*::FRT Δ*rplI*::*cat* | This study |
| Δ*rpmJ* Δ*prfC* | MG [illegible] | This study |
| Δ*rsmG* Δ*prfC* | MG Δ*rsmG*::FRT Δ*prfC*::*cat* | This study |
| Δ*tufA* Δ*prfC* | MG Δ*tufA*::FRT Δ*prfC*::*cat* | This study |
| Δ*rsxA* Δ*prfC* | MG Δ*rsxA*::FRT Δ*prfC*::*cat* | This study |
| Δ*yfjF* Δ*prfC* | MG Δ*prfC*::FRT Δ*yfjF*::*cat* | This study |

| **Plasmids** | **Description** | **Source** |
|---|---|---|
| pZS-P*tet*-m-y | Rep101; Amp$^r$ | (2) |
| pZS-P*tet*-m-TGA-y | Rep101; Amp$^r$ | (2) |
| pZS-P*tet*-eCFP-TGA-y | Rep101; Amp$^r$ | This study |
| pZS-P*tet*-lacZ | Rep101; Amp$^r$ | (2) |
| pEK4 | pMB1 ori; Amp$^r$ | (3) |
| pEK7 | pMB1 ori; Amp$^r$ | (3) |
| Dual-luc-15Tyr | pMB1 ori; Amp$^r$ | This study |
| pKD46 | Rep101; Amp$^r$ | (4) |
| pKD3 | R6K γ ori; Amp$^r$ and Cam$^r$ | (4) |
| pCP20 | Rep101(Ts); Amp$^r$ and Cam$^r$ | (4) |
| ASKA *E. coli* ORF library | pMB1 ori; Cam$^r$ | (5) |

| | | |
|---|---|---|
| pSZ-P*tet*-eCFP-TGA-y | Rep101; Amp$^r$ | This study |
| pKT tRNA$^{Trp}$ | pMB1 ori; Kan$^r$ | Lab collection |
| pSZ- P*tet*-mCherry-TGA-y-P*rrsC*-eCFP | Rep101; Amp$^r$ | This study |
| pSZ- P*tet*-mCherry-TGA-y-P*tnaC*-eCFP | Rep101; Amp$^r$ | This study |

**Table S3. Oligonucleotides used in this study.**

**References**


1. Baba, T., Ara, T., Hasegawa, M., Takai, Y., Okumura, Y., Baba, M., Datsenko, K.A., Tomita, M., Wanner, B.L. and Mori, H. (2006) Construction of *Escherichia coli* K-12 in-frame, single-gene knockout mutants: the Keio collection. *Molecular systems biology*, **2**, 2006.0008.
2. Fan, Y., Evans, C.R., Barber, K.W., Banerjee, K., Weiss, K.J., Margolin, W., Igoshin, O.A., Rinehart, J. and Ling, J. (2017) Heterogeneity of stop codon readthrough in single bacterial cells and implications for population fitness. *Molecular cell*, **67**, 826-836.
3. Kramer, E.B. and Farabaugh, P.J. (2007) The frequency of translational misreading errors in *E. coli* is largely determined by tRNA competition. *RNA*, **13**, 87-96.
4. Datsenko, K.A. and Wanner, B.L. (2000) One-step inactivation of chromosomal genes in *Escherichia coli* K-12 using PCR products. *Proc Natl Acad Sci U S A*, **97**, 6640-6645.
5. Kitagawa, M., Ara, T., Arifuzzaman, M., Ioka-Nakamichi, T., Inamoto, E., Toyonaga, H. and Mori, H. (2005) Complete set of ORF clones of *Escherichia coli* ASKA library (a complete set of *E. coli* K-12 ORF archive): unique resources for biological research. *DNA Res*, **12**, 291-299.
